# Environmentally sensitive hotspots in the methylome of the early human embryo

**DOI:** 10.1101/777508

**Authors:** Matt J. Silver, Ayden Saffari, Noah J. Kessler, Giriraj R. Chandak, Caroline H.D. Fall, Prachand Issarapu, Akshay Dedaniya, Modupeh Betts, Sophie E. Moore, Michael N. Routledge, Zdenko Herceg, Cyrille Cuenin, Maria Derakhshan, Philip T. James, David Monk, Andrew M. Prentice

**Affiliations:** Medical Research Council Unit The Gambia at the London School of Hygiene and Tropical Medicine, UK & Gambia; Department of Genetics, University of Cambridge, UK; Genomic Research on Complex Diseases (GRC Group), CSIR-Centre for Cellular and Molecular Biology, Hyderabad, India; MRC Lifecourse Epidemiology Unit, University of Southampton, Southampton General Hospital, UK; Department of Women and Children’s Health, King’s College London, UK; School of Medicine, University of Leeds, UK; School of Food and Biological Engineering, Jiangsu University, Zhenjiang, China; Epigenomics and Mechanisms Branch, International Agency for Research on Cancer, Lyon, France; Biomedical Research Centre, University of East Anglia, UK; Bellvitge Institute for Biomedical Research, Spain

## Abstract

In humans, DNA methylation marks inherited from gametes are largely erased following fertilisation, prior to construction of the embryonic methylome. Exploiting a natural experiment of seasonal variation including changes in diet and nutritional status in rural Gambia, we analysed two independent child cohorts and identified 259 CpGs showing consistent associations between season of conception (SoC) and DNA methylation. SoC effects were most apparent in early infancy, with evidence of attenuation by mid-childhood. SoC-associated CpGs were enriched for metastable epialleles, parent-of-origin specific methylation and germline DMRs, supporting a periconceptional environmental influence. Many SoC-sensitive CpGs overlapped enhancers or sites of active transcription in H1 ESCs and fetal tissues. Half were influenced but not determined by measured genetic variants that were independent of SoC. Environmental ‘hotspots’ providing a record of environmental influence at periconception constitute a valuable resource for investigating epigenetic mechanisms linking early exposures to lifelong health and disease.

## Introduction

DNA methylation (DNAm) plays an important role in a diverse range of epigenetically regulated processes in mammals including cell differentiation, X-chromosome inactivation, genomic imprinting and the silencing of transposable elements^1^. DNAm can influence gene expression and can in turn be influenced by molecular processes including differential action of methyltransferases and transcription factor binding^2,3^.

The human methylome is extensively remodelled in the very early embryo when parental gametic methylation marks are largely erased before acquisition of lineage and tissue-specific marks at implantation, gastrulation and beyond^4^. The days following conception may therefore offer a window of heightened sensitivity to external environmental influences, potentially stretching back to the period before conception coinciding with late maturation of oocytes and spermatozoa at loci that (partially) evade early embryonic reprogramming^5^.

The effects of early exposures on the mammalian methylome have been widely studied in animal models but multiple factors make this challenging in humans. Causal pathways are difficult to elucidate in observational studies, and even randomised experimental designs are prone to confounding due to reverse causation from exposure-related effects^6^.

Here we address these limitations by exploiting a natural experiment in rural Gambia where conceptions occur against a background of repeating annual patterns of dry (‘harvest’) and rainy (‘hungry’) seasons with accompanying significant changes in energy balance, diet composition, nutrient status and rates of infection^7,8^. We assess the influence of seasonality on DNAm in two Gambian child cohorts^9,10^, enabling robust identification of loci showing consistent effects at the ages of 24 months and 8-9 years (Fig. 1). Through prospective study designs we capture conceptions throughout the year and, in contrast to previous analyses in this population^11–13^, we use statistical models that make no prior assumptions about specific seasonal windows driving DNAm changes in offspring.

**Figure 1.**
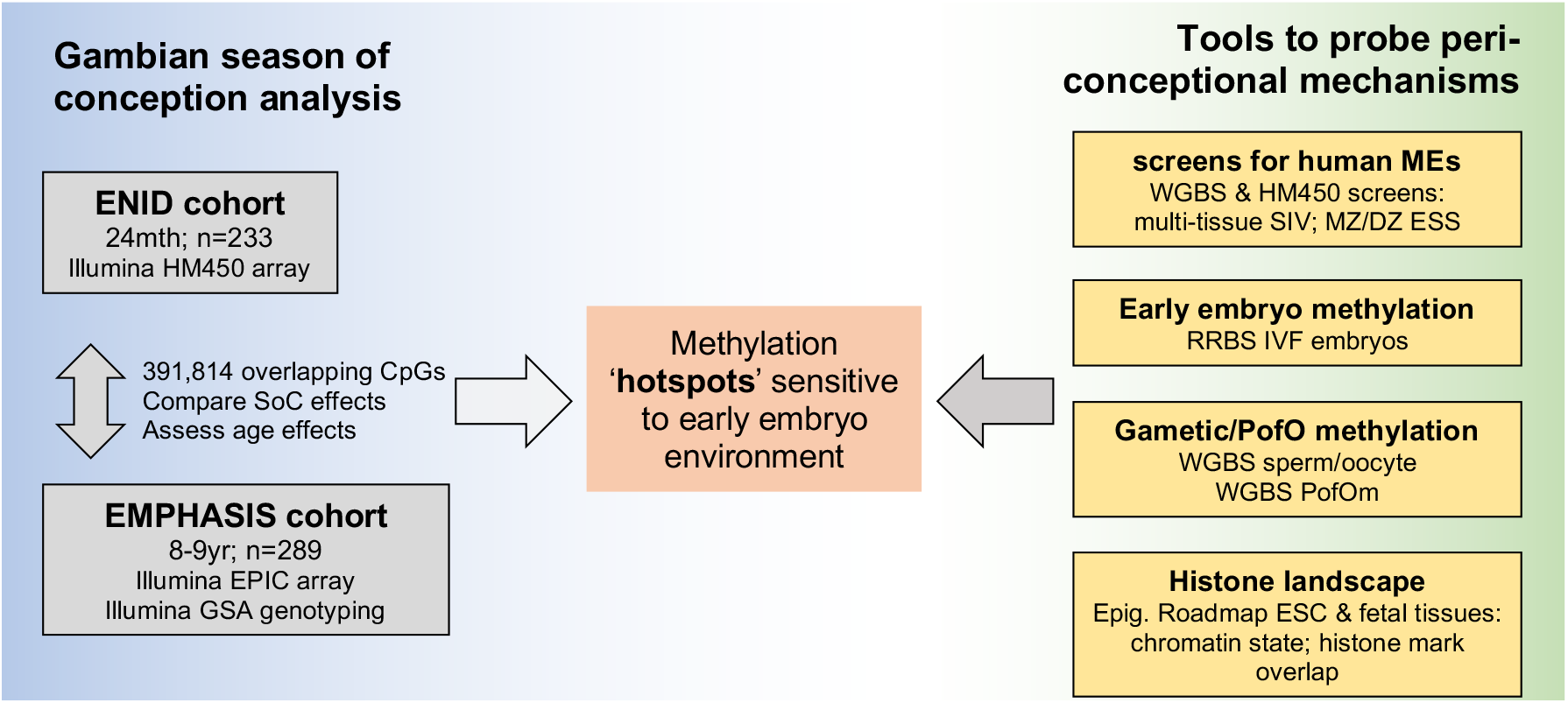
Study design. DNAm: DNA methylation; EPIC: Illumina Infinium MethylationEPIC BeadChip; Epig. Roadmap: Roadmap Epigenomics Consortium; ESC: embryonic stem cell; ESS: epigenetic supersimilarity; GSA: Global Screening Array; HM450: Illumina Infinium HumanMethylation450 BeadChip; IVF: in vitro fertilisation; MEs: metastable epialleles; MZ/DZ: monozygotic / dizygotic twins; PofO: parent of origin; RRBS: reduced representation bisulfite-seq; SIV: systemic interindividual variation; SoC: season of conception; WGBS: whole genome bisulfite-seq. See Tables 1 and 3 for further details of Gambian and public datasets used in this analysis.

We probe potential connections between season of conception (SoC)-associated loci and early embryonic events by leveraging published data on loci with evidence for the establishment of variable methylation states in the early embryo that persist in post-gastrulation and postnatal tissues; namely loci demonstrating systemic interindividual variation (SIV)^14,15^ and/or epigenetic supersimilarity (ESS)^15^ (Fig. 1). These loci bear the hallmarks of metastable epialleles (MEs), loci with methylation states that vary between individuals that were first identified in isogenic mice. MEs exhibit stable patterns of SIV indicating stochastic establishment of methylation marks prior to gastrulation when tissue differentiation begins^16^, and several MEs have been shown to be sensitive to periconceptional nutrition in mice^17^. These loci thus serve as useful tools for studying the effects of early environment on DNAm by enabling the use of accessible tissues (such as blood) that can serve as a proxy for systemic (cross-tissue) methylation, and by pinpointing the window of exposure to the periconceptional period^18^. We also investigate links with transposable elements and transcription factors associated with the establishment of methylation states in the early embryo, and assess the influence of genetic variation and gene-environment interactions. Finally, by comparing our results with public DNAm data obtained from sperm, oocytes and multi-stage human embryos, we investigate links between SoC-associated loci, histone marks, gametic and parent-of-origin specific methylation, and the establishment of DNAm states in early embryonic development.

Our identification of hotspots in the postnatal methylome that retain a record of environmental conditions during gametic maturation and/or in the very early embryo, provides a valuable resource for the investigation of epigenetic mechanisms linking early-life nutritional and other exposures to lifelong health and disease.

## Results

### Identification of Gambian season of conception associated CpGs

Key characteristics of the Gambian cohorts and samples analysed in this study are provided in Table 1. DNAm differences associated with season of conception are potentially confounded by season of sample collection effects in the ENID cohort (n=233) since samples were collected at age 24 months (Fig. 2A top). This is not the case in the older EMPHASIS cohort (n=289; age 8-9 yrs) where all samples were collected in the Gambian dry season (Fig. 2A bottom). To account for the potential influence of season of collection effects, we therefore compared year-round DNAm signatures across both cohorts by focussing on 391,814 autosomal CpGs (‘array background’) intersecting the Illumina HM450 and EPIC arrays used to measure DNAm in the ENID and EMPHASIS cohorts respectively (Table 2). We modelled the effect of date of conception on DNAm using Fourier (or ‘cosinor’) regression^19^ which makes no prior assumptions about specific seasonal windows that might drive DNAm changes in offspring (see Methods).

**Table 1.** Gambian seasonality-methylation analysis: cohort characteristics.

| Cohort | sample size | Age | % male | Tissue | methylation array |
| --- | --- | --- | --- | --- | --- |
| ENID<sup>9</sup> | 233 | 24mth | 50.6 | peripheral blood | Illumina Infinium HM450 |
| EMPHASIS<sup>10</sup> | 289 | 8-9y | 54.3 | peripheral blood | Illumina Infinium MethylationEPIC |

*Note: there is no overlap between individuals included in the two cohorts.*

**Table 2.** CpG sets considered in this analysis.

| CpG set | Number of CpGs | Notes |
| --- | --- | --- |
| Array background | 391,814 | Intersection of CpGs on Illumina HM450 (ENID) and EPIC (EMPHASIS) cohort arrays, post QC |
| SoC-associated CpGs | 768 | SoC-associated loci identified in the ENID (2yr) cohort (LRT FDR<5%) |
| **SoC-CpGs** | 259 | SoC-associated CpGs with SoC effect size (SoC methylation amplitude) > 4% |
| Matched controls | 259 | CpGs with similar methylation distributions to SoC-CpGs* |
| Random controls | 259 | Random sample from array background |

\*Matching methylation distributions determined by Kolmogorov-Smirnov tests (see Supplementary Fig. 15). QC: quality control; LRT: likelihood ratio test. See Methods for further details.

**Figure 2.**
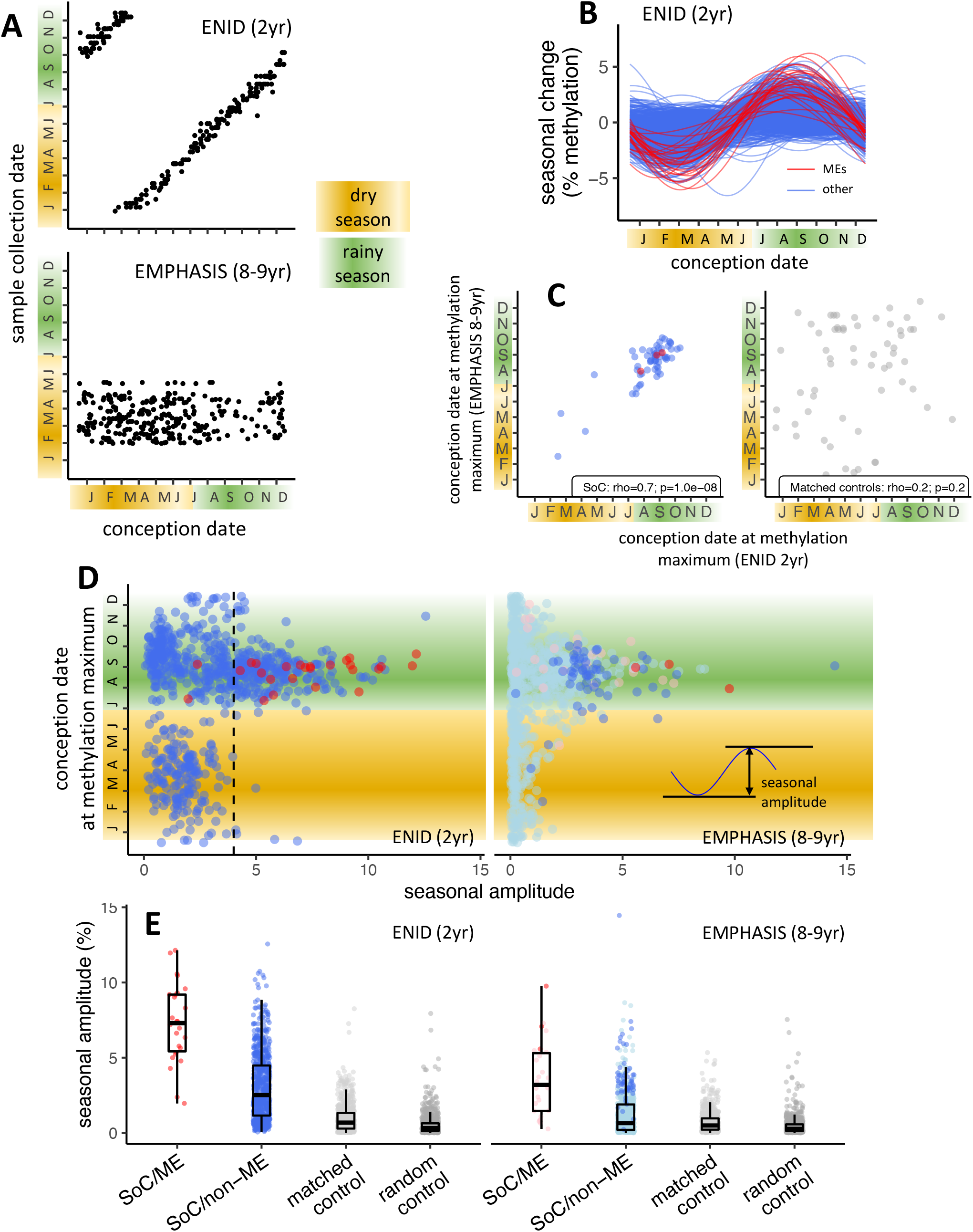
Identification of Gambian season of conception associated CpGs. **(A)** Relationship between date of conception and date of sample collection for ENID (top) and EMPHASIS (bottom) cohorts. **(B)** Modelled seasonal change in methylation for 768 SoC-associated loci (FDR<5%) in the ENID cohort. 26 ME CpGs are marked in red. **(C)** Conception date of modelled methylation maximum in each cohort for 61 CpGs significantly associated with SoC in both cohorts (left) and 61 matched controls (right). **(D)** (Left) Date of modelled DNAm maximum vs seasonal amplitude in each cohort for 768 CpGs significantly associated with SoC in the ENID cohort. MEs are marked in red. Dashed line indicates SoC amplitude threshold used to identify SoC-CpGs. (Right) same CpGs as left, but in the older EMPHASIS cohort. Significant SoC association for this cohort are marked in a darker colour. **(E)** (Left) Seasonal amplitudes for SoC-associated CpGs that are (red) or are not (blue) MEs; along with amplitudes for 768 matched and random controls (light/dark grey respectively). (Right) as left but in the older EMPHASIS cohort. For EMPHASIS significant SoC associations are marked in a darker colour. Boxes represent the middle 50% of the data (interquartile range, IQR); the line inside the box is the median, and whiskers represent values lying within 1.5 times the IQR.

We began by identifying 768 SoC-associated CpGs showing significant seasonal variation in 2-year olds from the ENID cohort with a false discovery rate (FDR)<5% (Table 2; Supplementary Table 1; Methods). Fourier regression models revealed a heterogeneous distribution of year-round methylation peaks and nadirs at these loci (Fig. 2B). SoC-associated loci were highly enriched for loci exhibiting SIV/ESS previously identified in multi-tissue screens in adult Caucasians^14,15^, hereafter named ‘MEs’ for short (Table 3; Fig. 2B, 26 ME CpGs marked in red; enrichment p = 2.5×10^−14^). More than twice as many of these loci were within 100bp of a putative ME (n=56; Supplementary Table 1). All identified loci showed increased seasonal amplitudes, defined as the distance between methylation peak and nadir, compared to matched and random controls (Supplementary Table 2; see Table 2 and Methods for justification and further details on selection of controls). Loci with the largest amplitudes tended to show increased methylation in conceptions in the Gambian rainy season (Fig. 2D left) in line with our previous observations in this population^11,12^.

**Table 3.**
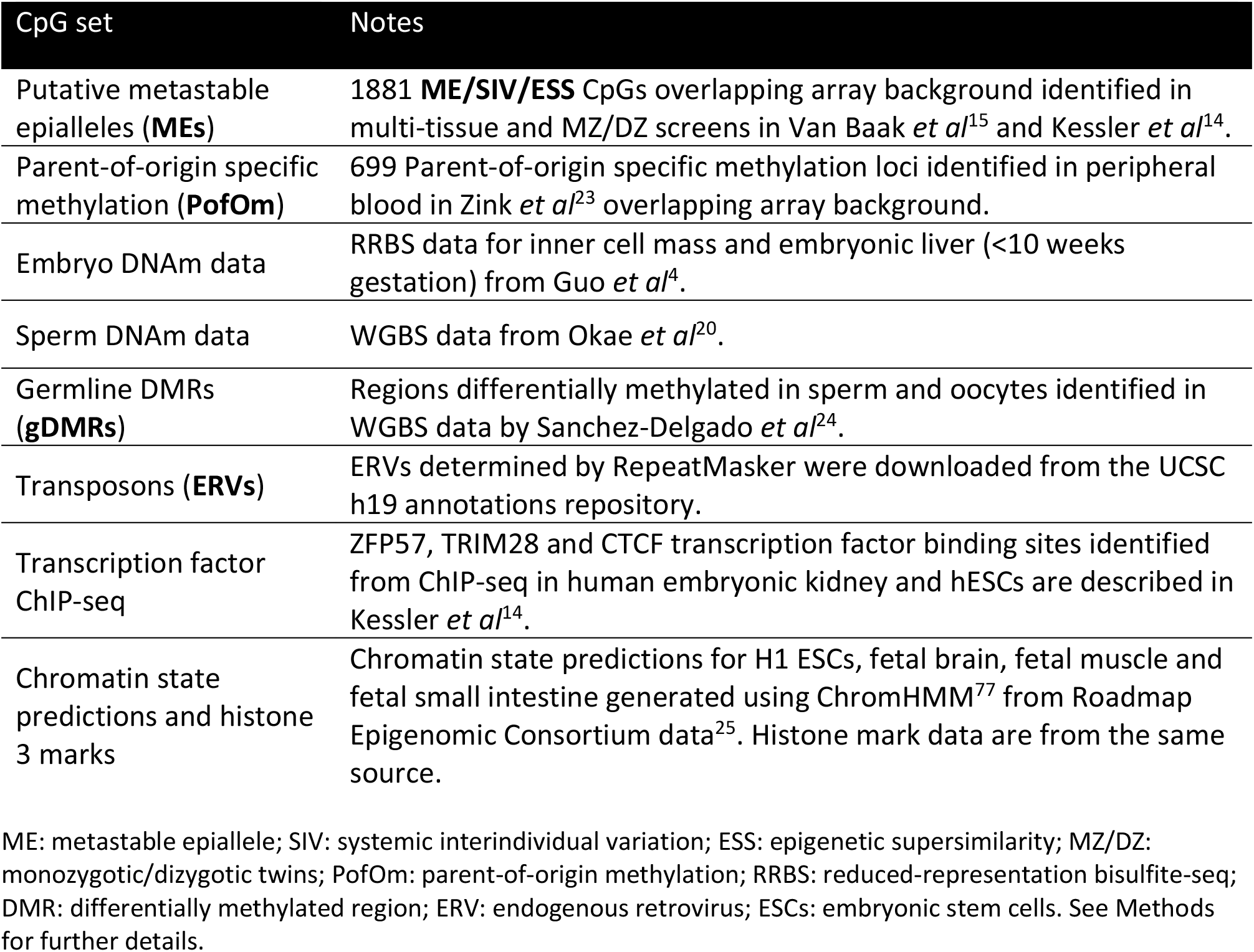
External datasets considered in this analysis.

Next, we analysed SoC effects at these 768 loci in 8-9-year olds from the EMPHASIS cohort. Mean methylation at SoC-associated loci was strongly correlated across cohorts (Supplementary Fig. 1), and we found evidence of a similar effect of increased methylation in conceptions in the Gambian rainy season (Fig. 2D right). 61 loci (including three ME CpGs) were also significantly associated with SoC (FDR<5%) in the older cohort (Fig. 2D right). Notably, the date of conception at methylation maximum was highly correlated across these two independent and different aged cohorts (Fig. 2C left; Spearman rho=0.7, p=1.0×10^−8^). No significant correlation was observed at matched controls with similar methylation distributions to SoC-associated loci (Fig. 2C right).

Given the strikingly similar seasonal patterns across the two cohorts, we next investigated reasons for the smaller number of SoC-associated loci (FDR<5%) in the older (EMPHASIS) cohort. Focussing on the 768 SoC-associated loci in the ENID cohort, we found significant and relatively large effect size (SoC amplitude) decreases in the older cohort, with SoC-effect attenuation most marked at loci that are also putative MEs (4.1% median methylation decrease, Wilcoxon p=2.8×10^−6^; non-ME CpGs: 1.9%, Wilcoxon p=1.5×10^−77;^ Fig. 2E; Supplementary Table 3). Corresponding SoC amplitude changes in matched and random controls were much smaller (0.2%, p=2.0×10^−8^ and 0.05%, p=0.02 respectively; Supplementary Table 3).

Our observation of a similar seasonal signature across two cohorts with different confounding structures (Figs. 2A, 2C & 2D), combined with evidence of SoC-effect attenuation in the older EMPHASIS cohort (Fig. 2E) led us to conclude that loci identified in the younger ENID cohort are robust sentinels of SoC-associated effects persisting at least until the age of 2 years.

Loci with larger SoC amplitudes showed a more consistent pattern of seasonal variation, both within and between cohorts (Figs. 2C & 2D). Reasoning that even transient molecular biomarkers of periconceptional environment in postnatal tissues could have biological significance, we therefore focussed on 259 SoC ‘hotspots’ or SoC-CpGs with FDR<5% and SoC amplitude ≥ 4% in the ENID cohort (Fig. 2D left, loci to the right of the dashed line; Supplementary Table 4).

SoC-CpGs are distributed throughout the genome and cluster together in several regions (Supplementary Fig. 2). Noting that the number of clusters is relatively insensitive to the inter-CpG distance used to define them (Supplementary Fig. 3), we identified 56 distinct SoC-CpG clusters and 105 ‘singletons’ (SoC-CpGs with no close neighbours) using a minimum inter-CpG distance of 5kbp (Supplementary Table 5). With this definition, 59% of SoC-CpGs fell within clusters (Supplementary Table 6). Of note, SoC effect amplitudes and cross-cohort correlations were greater at SoC-CpGs falling within clusters than with singletons (Supplementary Fig. 4).

Several SoC-CpG clusters extend over more than 500bp, notably a cluster mapping to *IGF1R* which spans 872bp and covers 7 CpGs (Supplementary Table 5, Supplementary Fig. 5). All but one of these 7 CpGs were also significantly associated with SoC (FDR<5%) in the older EMPHASIS cohort (Supplementary Table 4).

### Properties of SoC-CpGs

Compared to array background, SoC-CpGs are intermediately methylated, most notably at putative MEs (Fig. 3A), and they tend to fall within CpG islands compared to array background and matched controls (Fig. 3B).

**Figure 3.**
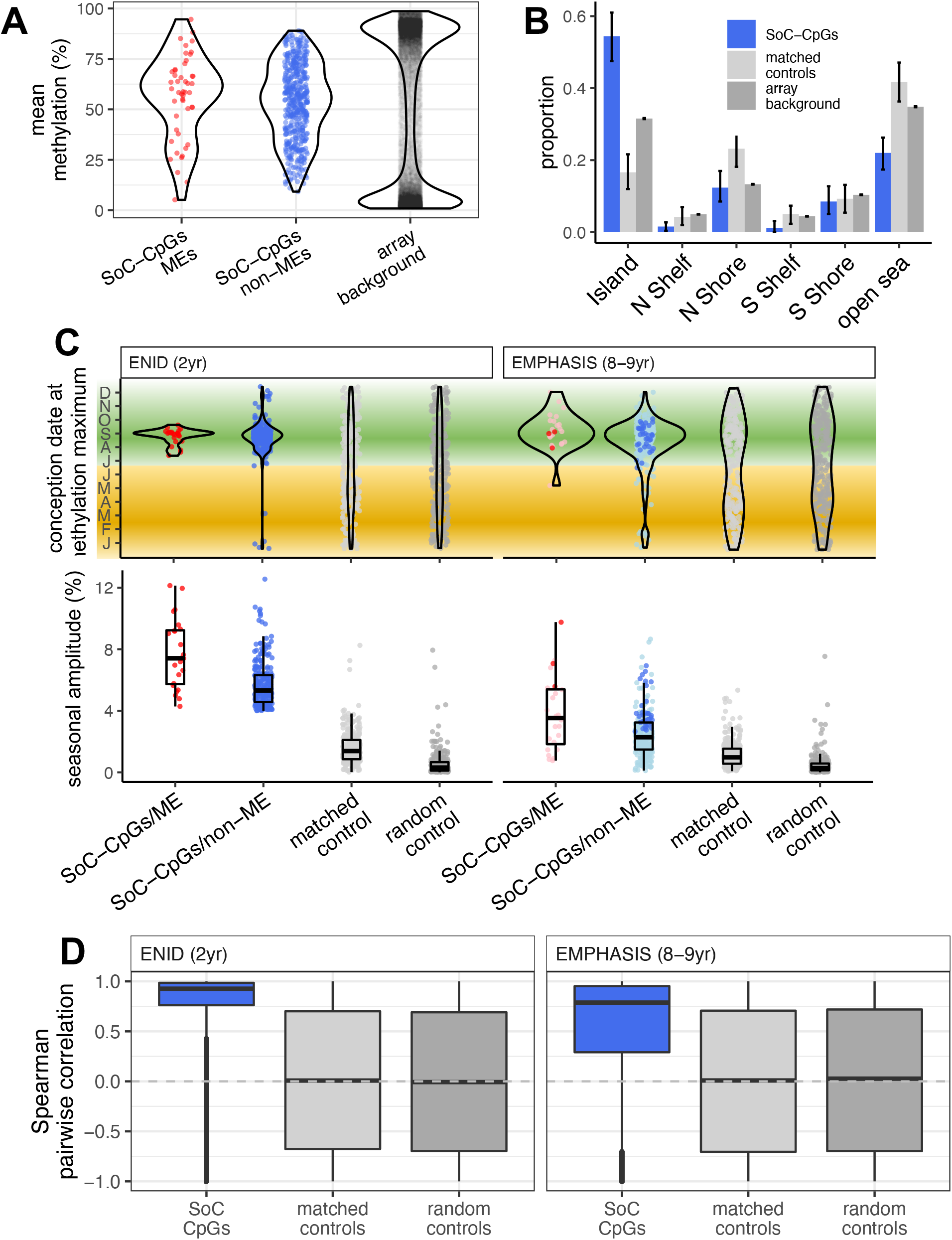
Properties of SoC-CpGs. **(A)** SoC-CpGs show increased intermediate methylation compared to array background (data from both cohorts combined). **(B)** Distribution of 259 SoC-CpGs (including MEs), 259 matched controls and array background with respect to CpG islands. Error bars are bootstrapped 95% CIs. N / S Shore / Shelf: North / South Shore / Shelf respectively (regions proximal to CpG Islands defined in Illumina manifest). **(C)** Date of conception at modelled methylation maxima (top) and seasonal amplitude (bottom) for 259 SoC-CpGs and 259 corresponding matched and random controls. Significant SoC associations (FDR<5%) in the EMPHASIS cohort are marked in a darker colour. Green and yellow bands indicate the extent of the rainy and dry seasons respectively. Boxplot elements as described in Fig. 2E. **(D)** Distribution of pairwise Spearman correlations for CpG sets in the ENID (left) and EMPHASIS (right) cohorts. Boxplot elements as described in Fig. 2E.

As anticipated (Fig. 2D), SoC-CpGs show a distinct pattern of methylation maxima for conceptions falling within the August-September period in both cohorts, most markedly at putative MEs with independent evidence of establishment in the early embryo (Fig. 3C top). The August to September period corresponds to the peak of the Gambian rainy season, a strong validation of our previous studies in babies and infants that focussed on conceptions at peak seasons only, with similar observations of higher methylation in conceptions at the peak of the Gambian rainy season compared to peak dry season^11–13,15^. Methylation minima fall within the February-April period, corresponding to the peak of the dry season (Supplementary Fig. 6). As expected (Fig. 2E), seasonal methylation amplitudes are larger at SoC-CpGs compared to controls and appear to decrease with age (Fig. 3C bottom; Supplementary Table 7).

Compared to array background, SoC-CpGs are highly enriched for MEs (21-fold enrichment, p=3.0×10^−23^, cluster-adjusted: 17-fold, p=3.1×10^−11^; Supplementary Table 8; see Methods for further details on cluster-based adjustments). The number of SoC-CpGs directly overlapping previously identified MEs is small (n=24), although a total of 49 SoC-CpGs (19%) fall within 100bp of a ME (Supplementary Table 4). Further investigation revealed that a large majority (n=19/24) of overlapping MEs were identified in our previous screen for SIV, with a smaller number exhibiting ESS (n=7/24; Supplementary Table 8). No MEs overlap methylation distribution-matched controls.

Finally, pairwise methylation states are highly correlated at a large majority of SoC-CpGs in both cohorts, so that the same individuals tend to have relatively high or low methylation at multiple SoC-CpGs (Fig. 3D). Pairwise correlations are not driven by increased correlation within SoC-CpG clusters (Supplementary Fig. 7), and methylation at matched and random controls is uncorrelated, thus reducing the possibility that these correlations are driven by statistical artefacts (Fig. 3D, Supplementary Fig. 7).

### Early stage embryo, gametic and parent-of-origin specific methylation

Given the strong enrichment for MEs within the set of SoC-CpGs, we next analysed links to methylation changes in early stage human embryos, as we have done previously for putative MEs identified in a whole-genome bisulfite-seq (WGBS) multi-tissue screen^14^. We aligned our data with public reduced representation bisulfite-seq (RRBS) data from human IVF embryos^4^ and obtained informative methylation calls for 112,380 array background CpGs covered at ≥ 10x read depth in inner cell mass (ICM, pre-gastrulation) and/or embryonic liver (post-gastrulation) tissues. As previously noted at putative MEs^14^, we found a distinctive pattern of increased incidence of intermediate methylation states at SoC-CpGs in post-gastrulation embryonic liver tissue, strongly contrasting with a general trend of genome-wide hyper- and hypo-methylation at loci mapping to array background (Fig. 4A). A similar pattern of increased incidence of intermediate methylation states was observed at distribution-matched controls.

**Figure 4.**
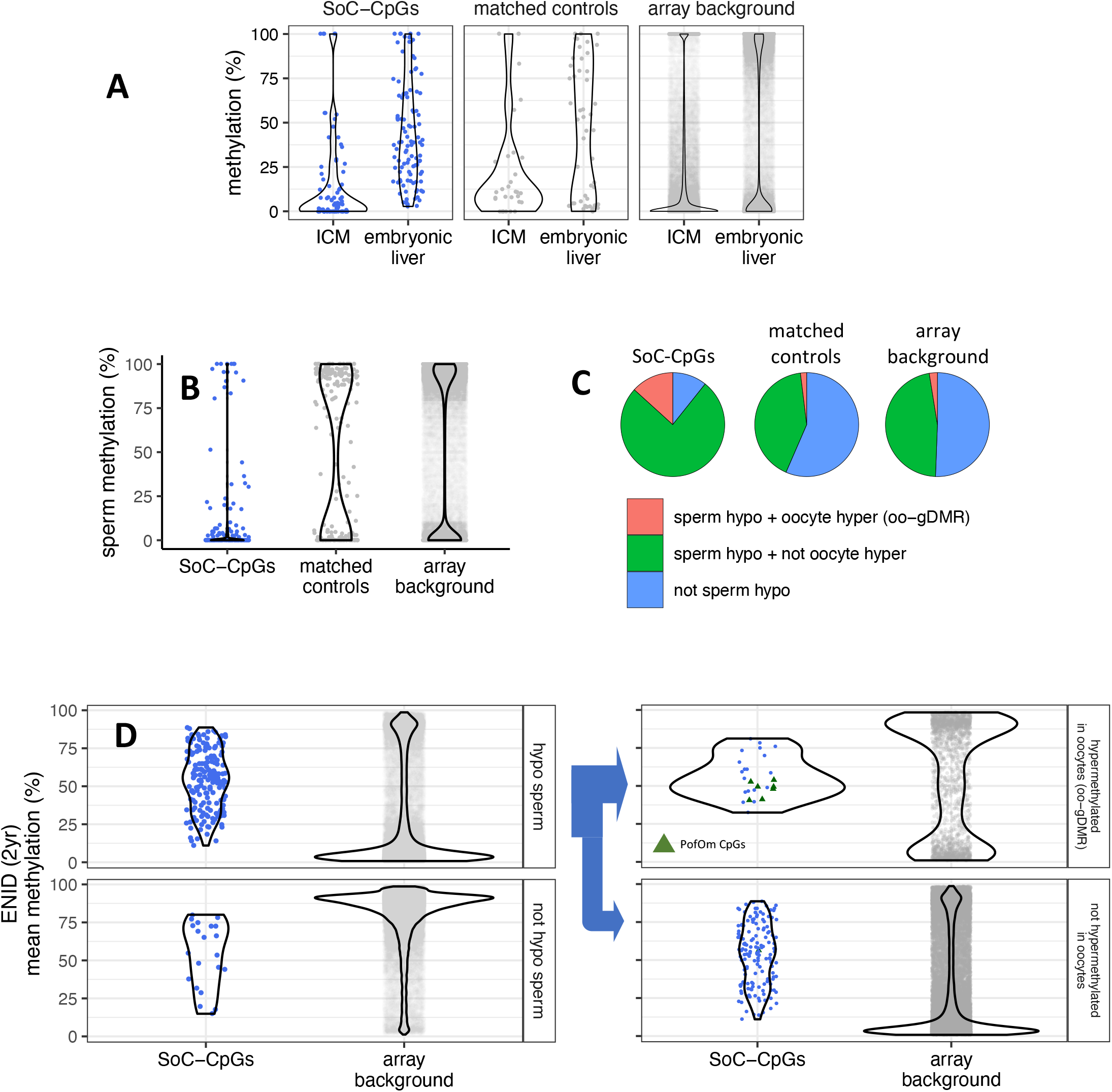
Early stage embryo, gametic and parent-of-origin specific methylation. **(A)** Methylation distribution of SoC-CpGs, matched controls and array background in pre-gastrulation inner cell mass (ICM) and post-gastrulation embryonic liver, measured in RRBS embryo methylation data from Guo *et al*^4^. Data comprises 112,380 CpGs covered at ≥10x in ICM and/or embryonic liver that overlap array background, including 118 SoC-CpGs and 51 matched controls. **(B)** Methylation distribution of SoC-CpGs, matched controls and array background in sperm WGBS data from from Okae *et al*^20^. Data comprises 294,240 CpGs covered at ≥ 10x including 196 SoC-CpGs and 207 matched controls. **(C)** Sperm methylation and oocyte gDMR (oo-gDMR) status at 196 SoC-CpGs covered at ≥ 10x in Okae *et al* sperm WGBS data. Sperm hypomethylation is defined as methylation ≤ 25%. oo-gDMRs defined as sperm methylation <25% and oocyte methylation >75% in WGBS analysis by Sanchez-Delgado. *et al*^24^. **(D)** Mean methylation at SoC-CpGs and array background measured in n=233 individuals in the ENID (2yr) cohort stratified by sperm and oocyte methylation status. Left: Methylation stratified according to sperm hypomethylation status (n=175 SoC-CpGs hypomethylated in sperm, n=21 not hypomethylated). Right: as left but with loci hypomethylated in sperm further stratified according to oo-gDMR status (n=26 SoC-CpGs hypermethylated in oocytes / oo-gDMR, n=149 not hypermethylated). Sperm / oo-gDMR status thresholds as for Fig. 4C. 8 PofOm CpGs (green triangles) are those identified in Zink *et al*^23^.

We previously observed consistent hypomethylation at MEs across all gametic and early embryonic developmental stages, most notably in sperm^14^. We tested the latter observation at SoC-CpGs by aligning our data with public sperm WGBS data^20^, restricting our analysis to 294,240 CpGs mapping to array background that were covered at ≥ 10x. SoC-CpGs tended to be hypomethylated in sperm, compared to loci mapping to matched control CpGs and array background respectively (Figs. 4B & 4C). Intermediate methylation states at SoC-CpGs were preserved in both Gambian cohorts irrespective of sperm methylation states, in contrast to array background CpGs where methylation distributions strongly reflected sperm hypomethylation status (ENID cohort: Fig. 4D left; EMPHASIS cohort Supplementary Fig. 8 left).

Our observation of an increased incidence of sperm hypomethylation at SoC-associated loci, together with existing evidence that imprinted genes may be sensitive to prenatal exposures^13,21,22^, prompted us to investigate a potential link between SoC-CpGs and parent-of-origin specific methylation (PofOm). A recent study used phased WGBS methylomes to identify regions of PofOm in 200 Icelanders^23^. We analysed 699 of these PofOm CpGs overlapping array background (Table 3) and observed strong enrichment for PofOm CpGs at SoC-CpGs and at all MEs on the array (18- and 15-fold enrichment, p=3.0×10^−8^ and 1.8×10^−36^ respectively; Supplementary Table 9; Fig. 4D right, PofOm CpGs marked as green triangles). No enrichment was observed at distribution-matched controls (Supplementary Table 9). PofOm enrichment at SoC-CpGs is driven by a large PofOm region spanning 6 CpGs on chr15 at *IGF1R* (Supplementary Table 4); along with two singleton PofOm SoC-CpGs, one on chr18 close to *PARD6G*, and the other in the Prader-Willi syndrome-associated imprinted region neighbouring *MAGEL2*, also on chr15. All of these loci have increased methylation in the rainy season, with SoC effect sizes (methylation amplitudes) ranging from 4.1-8.4% (median 6.1%; Supplementary Table 4).

Regions of PofOm detected in postnatal samples tend to be differentially methylated in gametes^23^, and may thus have evaded epigenetic reprogramming in the pre-implantation embryo^21^. We tested this directly by interrogating data from a whole-genome screen for germline differentially methylated regions (gDMRs) that persist to the blastocyst stage and beyond^24^. In this analysis, gDMRs were defined as contiguous 25-CpG regions that were hypomethylated (mean DNAm < 25%) in one gamete and hypermethylated (mean DNAm > 75%) in the other. We began by observing strong enrichment for oocyte (maternally methylated), but not sperm gDMRs, at all PofOm loci identified by Zink *et al*^23^ (Supplementary Table 9), confirming previous observations of an excess of PofOm loci that are methylated in oocytes only^23^. This enrichment was particularly strong for oocyte gDMRs (oo-gDMRs) persisting in placenta (Supplementary Table 9). We next analysed SoC-CpGs and MEs and again found evidence for strong enrichment of oocyte, but not sperm gDMRs at these loci (6.2-fold oo-gDMR enrichment, p=2.3×10^−16^ at SoC-CpGs; 2.9-fold, p=1.2×10^−24^ at MEs), including after adjustment for CpG clustering (Supplementary Table 9). Of note, 14% (36/259) of SoC-CpGs overlapped oo-gDMRs, in strong contrast to matched and random controls (Fig. 4C). These clustered into 19 distinct oo-gDMR regions (Supplementary Table 10) - more than six times the number identified as exhibiting PofOm by Zink *et al* (3 regions; Supplementary Table 4).

A large majority of SoC-CpGs that are hypomethylated in sperm are not oo-gDMRs (i.e. they are not hypermethylated in oocytes) (Fig. 4C & 4D bottom right), suggesting that factors associated with regional sperm hypomethylation rather than differential gametic methylation may be a key driver of sensitivity to periconceptional environment at these loci.

### SoC-CpG overlap with predicted chromatin states

We assessed the overlap of SoC-CpGs with predicted chromatin states generated from histone marks in various cell lines and tissues by the Roadmap Epigenomics Consortium^25^. Given our interest in methylation states associated with periconceptional environment that persist into early postnatal life, we focussed on data from H1 embryonic stem cells (ESCs) and fetal tissues (fetal brain, muscle and small intestine) derived from all 3 germ layers, described as having the ‘highest quality’ epigenomes (see Fig. 2 in Kundaje et al^25^). Around half of all SoC-CpGs overlapped sites with predicted transcriptional or regulatory function, with relatively few overlapping constitutive heterochromatic regions (Fig. 5). Overlaps with specific histone marks in H1 ESCs are given in Supplementary Fig. 9. As expected, given predicted chromatin states, many show a predominance of overlapping H3K4me1 and H3K27me3 marks and combinations thereof, suggestive of active or poised enhancers.

**Figure 5.**
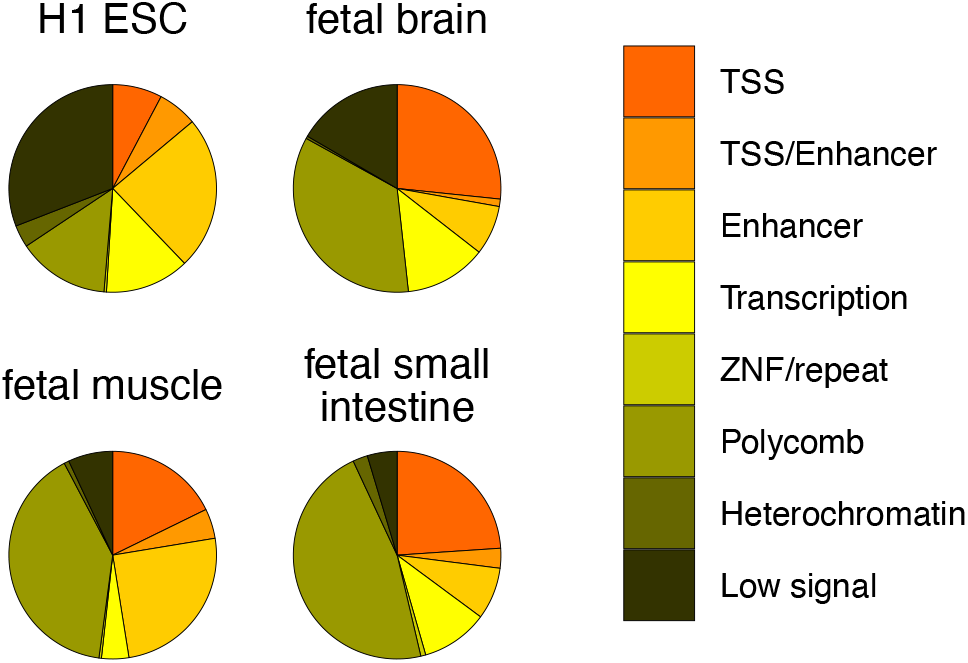
SoC-CpG overlap with predicted chromatin states. Chromatin states predicted by ChromHMM^77^ from chromatin marks in four cell lines and tissues generated by the Roadmap Epigenomics Consortium^25^. Predicted states for all 259 SoC-CpGs are shown. Predictions from the ChromHMM 15-state model are collapsed to 8 states for clarity. TSS: active transcription start site/flanking active TSS/bivalent or poisedTSS; TSS/Enhancer: flanking bivalent TSS/enhancer; Enhancer: Enhancer/bivalent enhancer/Genic enhancer; Transcription: Strong/weak transcription/ Transcription at gene 5’ and 3’; ZNF/repeat: zinc finger genes & repeats; Polycomb: Repressed/ weak repressed polycomb; Heterochromatin; Low signal: low signal in all marks states used as inputs to ChromHMM.

### Enrichment of transposable elements and transcription factors associated with genomic imprinting

Variable methylation states at MEs are associated with transposable elements (TEs) in murine models^26,27^, and we have previously observed enrichment for proximity to two classes of endogenous retroviruses, ERV1 and ERVK, at putative human MEs^13,14^. Here we found evidence that SoC-CpGs are enriched for proximity to ERV1 (Fig. 6A top) but not ERVK retroviral elements (Supplementary Fig. 10; Table 3). To maximise power, we tested the relationship between SoC amplitude and proximity to ERV1 in the extended set of 768 SoC-associated CpGs which includes loci with amplitudes <4% (Table 2). We found that SoC effect sizes were significantly larger at SoC-associated loci proximal to ERV1 transposons in both cohorts (Fig. 6B), but not at 768 controls with similar methylation distributions (Supplementary Fig. 11).

**Figure 6.**
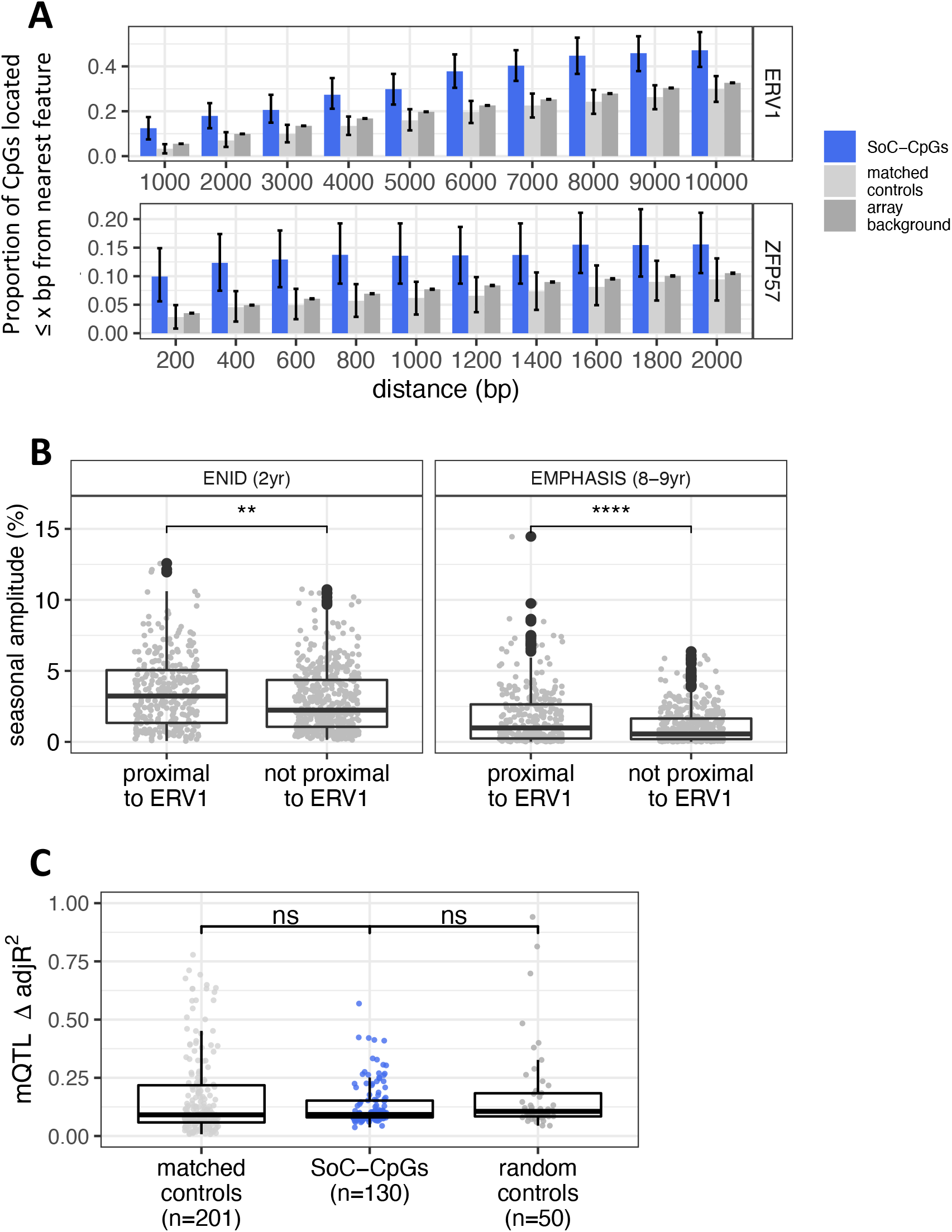
Links between endogenous retroviruses (ERVs), ZFP57 binding sites, genetic variation and DNAm at SoC-associated loci. **(A)** Proportion of SoC-CpGs, matched controls and array background CpGs proximal to ERV1 endogenous retroviral elements (top) and ZFP57 binding sites (bottom), within the specified distance. CpG clustering effects are removed by sampling a single CpG from each cluster (see Methods). Error bars are bootstrapped 95% CIs. **(B)** Difference in SoC effect size (seasonal amplitude) between SoC-associated loci proximal to ERV1 (≤10kbp; n=310) and not (>10kbp; n=458) in the ENID and EMPHASIS cohorts. Wilcoxon Rank Sum p-values under the null hypothesis of no difference between the two distributions tested: ENID p=0.002; EMPHASIS p=4.9×10^−5^. SoC-associated loci are those significantly associated with SoC in the ENID cohort (FDR<5%; n=768). Boxplot elements as described in Fig. 2E. **(C)** Proportion of methylation variance explained by mQTL for matched controls, SoC-CpGs and random controls. Only CpGs with at least one significant mQTL are plotted (n=201, 130 and 50 respectively; see Methods for further details). Boxplot elements as described in Fig. 2E.

Enrichment for PofOm and gDMRs at SoC-CpGs suggests a potential link to mechanisms implicated in the maintenance of PofOm and genomic imprinting in the early embryo. Our previous analysis of MEs identified from WGBS data found enrichment for proximal binding sites for 3 transcription factors (TFs: CTCF, ZFP57 and TRIM28) identified through ChIP-seq of embryonic stem and kidney cells that are linked to maintenance of PofOm at imprints^14^. Here we found evidence for enrichment of proximal ZFP57 binding sites within 2kbp of a SoC-CpG as previously observed at MEs, but we found no evidence for enrichment of proximal CTCF or TRIM28 binding sites in this array-based study (Fig. 6A bottom; Supplementary Fig. 10; Table 3).

### Influence of genotype

Genetic variation is a major driver of inter-individual variation in DNAm via methylation quantitative trait loci (mQTL)^28^. We explored the influence of mQTL on SoC-CpGs in the EMPHASIS cohort for which we had genotype data on 284 individuals measured at >2.6M SNPs after imputation from the Illumina Global Screening Array (GSA) and subsequent LD-pruning (see Methods). The majority of mQTL effects occur in *cis*^28^. In order to maximise power we therefore adopted a two-step approach where we performed separate screens for mQTLs in *cis* (defined as SNPs within 1Mb of an associated CpG^28^) and *trans* (all others), and compared our findings at SoC-CpGs with matched and random control CpGs (Methods). Half of SoC-CpGs had one or more associated mQTL compared with 78% and 19% of matched and random controls respectively. 92% of SoC-CpG mQTL were in *cis* (Table 4).

**Table 4.** Methylation quantitative trait loci (mQTL) associated with SoC-CpGs and controls.

| **CpG set** | **Number of CpGs with mQTL** | **Number of mQTL (*cis/trans*)** | **Median number of mQTL per CpG (IQR)** | **Methylation variance explained<sup>1</sup>** |
| --- | --- | --- | --- | --- |
| **SoC-CpGs** | 130 (50%) | 2771 (2549/222) | 6 (2-30) | 0.09 (0.08-0.15) |
| **Matched controls** | 201 (78%) | 7886 (7417/469) | 15 (4-50) | 0.09 (0.06-0.21) |
| **Random controls** | 50 (19%) | 1512 (1476/36) | 7 (2-35) | 0.1 (0.08-0.18) |

<sup>1</sup> delta adjusted R<sup>2</sup> (see Methods)

We next compared methylation variance explained by significant mQTL using adjusted R^2^ values for all SoC-CpGs and controls with at least one genome wide significant mQTL (FDR<5%; n=130, 201 and 50 CpGs for SoC-CpGs, matched and random controls respectively; Table 4). These values were compared to a baseline model that included the same set of covariates (principal components, age and sex) used in Fourier regression models for the main seasonality analysis, in order to account for potential differences in additional variance explained by other covariates and unmeasured factors (see Methods). There was no difference in additional variance explained by significant mQTL between SoC-CpGs and both sets of control CpGs (Fig. 6C; Table 4).

To assess the potential for genetic confounding of SoC-associated DNAm signals at SoC-CpGs with associated mQTL, we tested all SoC-CpG-mQTL for association with season of conception using an allelic model. After accounting for multiple testing, no significant SoC-mQTL associations were identified (Supplementary Table 11 and Methods). Our observations that i) SoC-CpGs are distributed throughout the genome; ii) SoC-CpG mQTL occur primarily in *cis*; and iii) none are associated with season of conception; strongly suggest that SoC-methylation associations at SoC-CpGs are not confounded by genetic variation.

Finally, we searched for gene-environment (SoC) interaction (GxE) effects, again performing separate tests for SNPs in *cis* and *trans.* No GxE associations were identified after correcting for multiple testing (see Methods).

### Influence of genetic ancestry

Eighty percent of the population of the Kiang West region of The Gambia from which the cohorts analysed in this study are drawn are of Mandinka ethnicity, with the majority of the remainder Fula^29^. This is evident from a genome-wide principal component analysis (PCA) of genetic variation in the EMPHASIS cohort, where we observed a distinct cluster of 16 individuals from a single village which is predominantly Fula (Supplementary Fig. 12). Individuals in the main ENID cohort were drawn from the same Kiang West villages as the EMPHASIS study, but we were unable to directly adjust for potential confounding effects due to genetic ancestry since no genetic data was available for this cohort. Based upon the EMPHASIS cohort PCA and our knowledge of village population structures, we reasoned that village of origin is a useful proxy for genetic ancestry in the ENID cohort and performed a sensitivity analysis with an additional covariate dichotomised according to whether an individual came from the predominantly Fula village. The first two genetic principal components were used as adjustment covariates for the corresponding EMPHASIS analysis. Results from this ethnicity-adjusted sensitivity analysis were not materially different from those obtained for the main analysis (Supplementary Fig. 13).

### Overlap of SoC-CpGs with existing studies

To place our findings in the context of existing literature on associations between DNAm and nutrition-related exposures, including exposure to famine conditions in gestation and previous reported associations with Gambian SoC, we checked for overlaps between SoC-CpGs and loci identified in a recent review by James et al^22^. Many cited studies including the majority of previous work in The Gambia used pyrosequencing and other methylation platforms targeting loci not covered by Illumina arrays. However, a total of 57 previously identified loci did overlap or partially overlap array background. None of these overlapped a SoC-CpG within 1kbp. We also checked for overlaps with the larger set of SoC-associated CpGs not passing the 4% minimum effect size threshold, and found a single CpG (cg17434309) mapping to *IGF2* that was within 1kbp of two previously identified loci, one linking maternal plasma vitamin B12 with cord blood methylation^30^, and the second linking gestational famine to blood methylation in older adults^31^.

We next looked for overlaps between SoC-CpGs and CpGs identified in the EWAS Catalog (http://ewascatalog.org/), a manually curated database of significant results (p<1×10^−4^) from published epigenome-wide association studies (EWAS). This search produced published associations for 167 out of the 259 SoC-CpGs, mapping to 27 unique traits covering a range of pre- and post-natal exposures (Supplementary Table 12). Noteworthy amongst the most frequently reported associations with SoC-CpGs (Supplementary Table 13) in the context of our study were those with sex, gestational age, maternal smoking in pregnancy, maternal plasma folate levels and adult body mass index (BMI) with 109, 45, 16, 6 and 1 associated SoC-CpG(s) respectively.

We investigated some of these links using data from the ENID cohort considered in our main analysis and confirmed multiple significant associations with infant sex but not gestational age or maternal folate at conception (Supplementary Table 14). Links with adult BMI and maternal smoking were not considered as adult BMI was not available and the incidence of smoking is extremely low in our study population.

All Fourier regression models in our main season of conception analysis included sex as an adjustment covariate. The finding that multiple SoC-CpGs were associated with sex in the EWAS Catalog, with replication of this association in the ENID cohort, was therefore surprising. This prompted us to check for a residual confounding effect due to sex by repeating our analysis with methylation values pre-adjusted for sex, prior to running the full regression models, with and without further adjustment for sex. This produced near identical results to the main analysis without pre-adjustment, suggesting that the observed SoC associations were not driven by confounding due to sex.

Finally we searched for SoC-CpGs within 1kbp of SNP associations (p<1×10^−5^) in the GWAS Catalog^32^, since DNAm could mediate GWAS signals in genomic regions where functional effects are difficult to elucidate^33^. 11 SoC-CpGs mapped to a total of 12 SNPs associated with 8 unique traits in the GWAS Catalog (Supplementary Table 15). Notable traits from a developmental programming perspective were those linked to childhood obesity^34^ (2 SoC-CpGs) and QRS traits associated with cardiovascular mortality in adults (1 SoC-CpG)^35,36^.

## Discussion

We have exploited a natural, seasonal experiment in rural Gambia whereby human conceptions are ‘randomised’ to contrasting environmental (especially dietary) conditions to examine whether this differential exposure leaves a discernible signature on the offspring methylome. We analysed methylation data from two independent, different-aged cohorts and identified 259 ‘SoC-CpGs’ with evidence of sensitivity to season of conception (SoC) in infancy. We found evidence of SoC effect attenuation in the older cohort, but SoC-related temporal patterns were nonetheless strikingly similar, suggestive of a common effect of periconceptional environment. Importantly, these cohorts have contrasting confounding structures, notably with regard to the timing of sample collection; the latter eliminating potential confounding due to seasonal differences in leukocyte composition. These results, derived from analysis of Illumina array data with limited coverage, suggest there may be many more hotspots sensitive to the periconceptional environment across the human methylome.

This analysis builds on previous epigenetic studies in this setting that have focussed on single cohorts and analysed methylation differences between individuals conceived at the peaks of the Gambian dry and rainy seasons only^11–13,15,37^. Increased methylation in offspring conceived at the peak of the Gambian rainy season is consistent with previous findings and this observation is now greatly strengthened by the application of Fourier regression to model year-round conceptions – an approach that makes no prior assumption of when methylation peaks and nadirs may occur. The number of identified SoC-CpGs is also substantially increased in this study. Comparisons with array background and control CpGs matching SoC-CpG methylation distributions increase confidence that these findings are not statistical artefacts.

Triangulation with other public data provides multiple lines of evidence supporting the notion that methylation states at these loci are established in the periconceptional period. First, they are highly enriched for putative MEs and related loci identified in other studies with characteristic methylation signatures suggestive of establishment early in embryonic development^14,15^. Second, like MEs, season-associated loci exhibit unusual methylation dynamics in early stage embryos^14^. Third, they have distinctive gametic methylation patterns, notably hypomethylation in sperm (in common with putative MEs^14^), and differential gametic and parent-of-origin specific methylation in a subset. Increased incidence of hypomethylated states in sperm at SoC-CpGs may reflect their enrichment at CpG islands^38^, sequence features that are largely refractory to protamine exchange, with the possibility for retaining epigenetic function associated with histone modifications into the early embryo^39^. Fourth, many overlap H3K4me1 and H3K27me3 marks which coordinate transient gene expression and early lineage commitment in human ESCs, in part through demarking poised enhancers^40^.

A large majority of SoC-CpGs have not previously been identified as MEs, but given the supporting evidence described above, we speculate that many are likely to be so. Indeed, evidence of attenuation of SoC effects with age suggests that, to the extent that interindividual variation is driven by periconceptional environmental factors, screens for putative MEs (including ESS and SIV) in adult tissues used as a reference in this analysis may be missing metastable regions which are more pronounced earlier in the life course. Evidence of much larger SoC effect sizes at known MEs in both Gambian cohorts supports this.

The observed effect attenuation in the older (8-9yr) cohort also has implications for detecting the effect of periconceptional exposures on DNAm in samples collected beyond the neonatal and early childhood periods, an important consideration for epigenetic epidemiological studies since non-persisting methylation differences could still have a significant impact on early developmental trajectories with life-long consequences^41,42^.

Intra-individual methylation states at SoC-associated loci are highly correlated across loci despite being distributed throughout the genome, suggesting that a common mechanism is at play. This contrasts with a recent study of murine MEs located within intracisternal A particle insertions (IAPs, of which the *Agouti* locus is a paradigm example), where no intra-individual correlation between stochastic methylation states was observed, although the mice were not exposed to different environments^27^.

Potential insights into mechanisms linking periconceptional environment to DNAm changes in postnatal tissues come from our investigations of the methylation status and genomic context of SoC-CpGs.

First, we observed strong enrichment for germline differentially methylated regions, with 14% of SoC-CpGs overlapping 19 gDMRs hypomethylated in sperm and hypermethylated in oocytes. A minority of these show evidence of PofOm persisting in postnatal tissues. This observation aligns with a growing body of evidence linking early environment, notably nutritional factors involved in one-carbon (C1) metabolism, with methylation at imprinted regions^21,22^. Indeed we have previously noted an association between season of conception and several C1 metabolites at a maternally imprinted region at the small non-coding RNA *VTRNA2-1*^13^, consistent with evidence of ‘polymorphic imprinting’ linked to prenatal environment at this locus^23,43^. Furthermore, we previously found strong enrichment for proximal binding sites of several transcription factors (TFs) associated with the maintenance of PofOm in the early embryo at MEs detected in a WGBS screen^14^. We were only able to replicate enrichment for one of these, ZFP57, at SoC-CpGs in this study. This may reflect the relatively small proportion of PofOm loci in the set of SoC-CpGs, or factors related to the limited methylome coverage of Illumina arrays. More targeted experimental work is required to determine the extent of SoC effects at imprinted loci, especially given our observation that SoC-CpGs are often proximal to ERV transposable elements that have recently been shown to drive the establishment of germline-derived maternal PofOm^44^. Hotspots with evidence of PofOm could be driven by an environmentally sensitive gain of methylation on the paternal allele that is propagated through development, incomplete reprogramming on the maternal allele leaving residual traces of methylated cytosines, or modest *de novo* methylation at some later point. A deeper understanding of mechanisms will require further investigation in cell and animal models.

Second, our observation of enrichment for proximity to ERV1 transposable elements at SoC-CpGs aligns with our previous finding at MEs^13,14^, and is notable since most environment-sensitive mouse MEs are associated with IAPs (which are rodent-specific ERVs)^27^. KRAB zinc-finger protein (KZFP)-mediated repression of transposable elements (TEs) including ERVs has also been proposed as a driver of the rapid evolution of gene regulation^45^. The KZFP ZFP57 is particularly interesting in this respect since its binding to DNA is linked both to repression of TEs and to the maintenance of genomic imprints in the pre-implantation embryo^46,47^. We previously identified a putative SoC-associated DMR in the *ZFP57* promoter in blood from younger Gambian infants (mean age 3.6 months)^13^. It is possible that non-replication of the SoC-association at *ZFP57* in the older Gambian cohort (and of the *VTRNA2-1* SoC-DMR mentioned above which was also identified in younger infants) reflects the more general attenuation of SoC effects described above. Interestingly there is some evidence that the *ZFP57* DMR, which lies 3kb upstream of the transcription start site, is established in the early embryo^15,48^, and that DNAm at this locus measured in neonatal blood is associated with maternal folate levels^49–51^. Given the important function of ZFP57 in pre-implantation methylation dynamics, its potential role as an environmentally sensitive regulator of multi-locus DNAm effects remains an open question.

Third, DNAm at SoC-CpGs is enriched for intermediate methylation states. Intermediate methylation has also been observed at MEs in Gambians and in non-Africans^11,12,37,52^, and this coincides with a similar observation at MEs in post-gastrulation embryonic tissues^14^. Intermediate methylation appears to be predominantly driven by variegated (intercellular) differences in methylation status within a sampled tissue, rather than by allele-specific methylation^53^. Observed differences in aggregate methylation at SoC-CpGs could thus reflect a direct influence of periconceptional environment on the establishment and maintenance of DNAm states in the early embryo at the cellular level, or ‘epigenetic selection’ whereby epigenetically distinct cells form a substrate for clonal selection during development, for example as a potential adaptation to differential metabolic exposures^54^. Stronger enrichment for putative MEs exhibiting SIV compared to those identified through ESS supports establishment in the post-implantation embryo, since methylation differences at ESS loci are presumed to originate in the pre-implantation embryo prior to embryo cleavage in MZ twins^15^.

The largest SoC-CpG cluster with 7 CpGs is on chromosome 15 and falls within the second intron of *IGF1R.* This bears the hallmarks of being a promoter or active/poised enhancer in multiple cell lines (Supplementary Fig. 14). Six CpGs in the *IGF1R* cluster have evidence of PofOm and show a relatively large SoC effect size (median SoC methylation amplitude) of 6.1%. Zink *et al*^23^ were unable to demonstrate PofO allele-specific expression (PofO-ASE) in this region although others have found evidence of maternal imprinting of an intronic lncRNA at this gene in cancerous cells^55,56^. Loss of IGF1 receptors gives rise to a major decrease in expression at multiple imprinted genes in mice, suggesting a pathway by which *IGF1R* might regulate growth and metabolism during early development^57^. IGF1R signalling is implicated in fetal growth, glucose metabolism and cancer^58–60^, and DNAm differences at *IGF1R* have been observed in birthweight-discordant adult twins^61^. Another SoC-CpG with evidence of PofOm, also on chromosome 15, falls within the known Prader-Willi syndrome-associated paternally expressed imprinted region.

Our observation that 109 out of 259 SoC-CpGs have been associated with sex in previous studies is intriguing. Given the relatively small number (compared to array size) of autosomal sex-linked loci identified in large studies on the Illumina 450k array^62,63^, this represents a very strong enrichment. We replicated a significant sex association at these loci in the ENID cohort analysed in this study. Our regression analyses were adjusted for sex, and additional sensitivity analyses with DNAm pre-adjusted for sex strongly suggest that our main findings are not confounded by sex. Interestingly, sex-associated loci are enriched at imprinted regions and sex-discordance at autosomal CpGs has been linked to androgen exposures *in utero*^62,64^. There is also evidence of sex differences in methylation at *DNMT3A/B* and *TET1* genes involved in *de novo* methylation and de-methylation pathways^62,64^, suggesting a possible interaction between sex-linked epigenetic changes and periconceptional environment during reprogramming in the early embryo.

DNAm is influenced by genotype and the latter is therefore a potential confounder when studying the effects of environmental exposures in human populations. A strength of our quasi-randomised Gambian seasonal model is that it minimises the potential for genetic confounding of modelled seasonal DNAm patterns, on the assumption that the timing of conceptions is not linked to genetic variants influencing DNAm. However, it is still possible that such variants might confound our observations, for example if they promote embryo survival under conditions of environmental stress. We tested this possibility using genetic data available for the EMPHASIS cohort and found no evidence of SoC-associated genetic variants driving inter-individual methylation differences at SoC-associated loci in *cis* or *trans*.

However, we did find that half of SoC-CpGs had at least one associated mQTL, indicating the presence of independent additive effects of environment and genetics at these loci, as has been suggested previously at other loci sensitive to pre/periconceptional nutritional exposures^31,65^. We have previously argued that the definition of MEs should be extended to include genomic regions whose DNAm state is under partial but non-deterministic genetic influence in genetically heterogeneous human populations^14^, and we would argue that the above observations at SoC-CpGs that exhibit many of the characteristics of MEs support this. Further analysis in larger datasets with genome sequencing data combined with functional analysis using cell models will be required to fully understand the relative contributions of environment and genetics to DNAm variation at regions of the type highlighted in this study.

Further work is required to investigate the functional relevance of DNAm changes at SoC-CpGs, some of which are relatively small (around 4% SoC amplitude). However, we can speculate that observed methylation changes, which may reflect changes in the chromatin landscape, have the potential to influence gene expression and early development, since our chromatin state analysis predicts that many overlap regions with functional significance in H1 ESCs and fetal tissues. A similarly modest DNAm change at a locus in the *POMC* gene that is associated with SoC in blood in younger Gambian infants has been linked to obesity risk in German children and adults^37,66^, and to differential transcription factor binding and differences in *POMC* expression^66^.

We have previously shown that several metabolites involved in one-carbon methylation pathways show significant seasonal variation in maternal blood plasma in this population which is largely dependent on subsistence farming^8,12^. While we suspect that seasonal differences in nutrition are likely drivers of the effects that we have observed, analysis of the links between maternal nutritional biomarkers and DNAm is challenging due to the complex interdependence of one-carbon metabolites acting as substrates and cofactors within one-carbon pathways^67^. Furthermore, it is possible that DNAm differences are linked to other aspects of Gambian seasonality, such as variation in pesticide use or infectious disease burden^7,68^.

Limitations of this analysis include a lack of longitudinal methylation data that would allow us to directly test SoC effect attenuation in individuals from a single cohort. We also lack genetic data for the ENID cohort, so that our genetic analyses are restricted to the older EMPHASIS cohort where SoC effects are smaller. Furthermore, comparator data on MEs and gamete, embryo and PofO methylation are derived from non-African individuals so we cannot assess the influence of ethnicity on these analyses.

There is increasing interest in the phenomenon of methylation variability as a marker of disease and of prenatal adversity^54,69^, and in genetic variation as a potential driver of methylation variance^70^. This raises the possibility that certain genetic variants could have been selected through their ability to enable graded, environmentally-responsive methylation patterns at MEs and SoC-associated loci that are able to sense the periconceptional environment, record the information, and adapt the phenotype accordingly. Our gene-environment interaction analysis likely lacked power to detect gene-environment (SoC) interactions, so that it was not possible to investigate the relative contributions of stochastic, environmental and genetically-mediated variance effects on the establishment of periconceptional SoC-associated methylation states in this study. Nonetheless, the proposition that environmentally sensitive epigenetic signals are selected through their ability to direct phenotypic development to better fit the anticipated future environment, leading to maladaptation and future disease if the environment changes, is intriguing and worthy of further investigation^5^.

## Materials and Methods

### Gambian cohorts and sample processing

Detailed descriptions of the Gambian cohorts analysed in the season of conception study are published elsewhere^9,10^. Briefly, for the younger cohort, blood samples from 233 children aged 24 months (median[IQR]: 731[729,733] days old) were collected from participants in the **E**arly **N**utrition and **I**mmune **D**evelopment (“ENID”) study^9^. DNA was extracted, bisulfite-converted and hybridised to Illumina HumanMethylation450 (hereafter “HM450”) arrays following standard protocols (see Van Baak *et al*^15^ for further details). For the older cohort, DNA was extracted from blood samples from 289 Gambian children aged 8-9 (9.0 [8.6,9.2] years) participating in the **E**pigenetic **M**echanisms linking **P**re-conceptional nutrition and **H**ealth **As**sessed in **I**ndia and **S**ub-Saharan Africa (“EMPHASIS”) study^10^, and was bisulfite-converted and hybridised to Illumina Infinium Methylation EPIC (hereafter “EPIC”) arrays, again using standard protocols.

For the ENID cohort, date of conception was calculated from fetal gestational age estimates obtained by ultrasound at the mother’s first ‘booking’ appointment. The same method was used for the EMPHASIS cohort, except for n=71 pregnancies that were > 24 weeks gestation at booking meaning that GA could not be accurately determined by ultrasound^10,71^. In this case date of conception was calculated as date of birth minus 280 days which is the average gestational length for this population.

### Methylation array pre-processing and normalisation

Raw intensity IDAT files from the HM450 and EPIC arrays were processed using the *meffil*^72^ package in R (v3.6.1) using standard *meffil* defaults. Briefly, this comprised probe and sample quality control steps (filtering on bisulfite conversion efficiency, low probe detection p-values and bead numbers, high number of failed samples per probe, high number of failed probes per sample); methylation-derived sex checks; removal of ambiguously mapping (i.e. cross-hybridising) probes^73^; removal of probes containing SNPs at the CpG site or at a single base extension; and removal of non-autosomal CpGs. Following filtering, methylation data was normalised with dye-bias and background correction using the *noob* method, followed by *Functional Normalisation* to reduce technical variation based on principal component analysis of control probes on the arrays^74^. After pre-processing and normalisation, methylation data comprised methylation Beta values for 421,026 CpGs on the HM450 array for 233 individuals from the ENID cohort, and 802,283 CpGs on the EPIC array for 289 individuals from the EMPHASIS cohort. Finally, 391,814 CpGs intersecting both arrays were carried forward for statistical analysis.

### Statistical modelling

Variation of DNAm with date of conception was modelled using Fourier regression^19^. This models the relationship between a response variable (here DNAm) and a cyclical predictor (date of conception). The effect of the latter is assumed to be cyclical due to annually varying seasonality patterns, so that the modelled effect for an individual conceived on the 31^st^ December should be ‘close’ to that for an individual conceived on the 1^st^ of January. This is achieved by deconvolving the conception date (predictor) into a series of pairs of sin and cosine terms, and obtaining estimates for the regression coefficients β and γ in the following model:

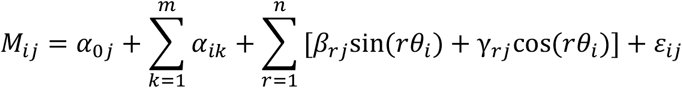

Where, for individual *i* and CpG *j*:

*M_ij_* is the logit-transformed methylation Beta value^75^;

*α_0j_* is an intercept term;

*α_ik_* is the *k^th^* of *m* adjustment covariates;

*θ_i_* is the date of conception in radians in the interval [0, 2π], with 1^st^ January = 0 and 31^st^ December = 2π, modelled as *n* pairs of Fourier terms, sin *θ_i_* + cos *θ_i_* + … + sin *n*θ_i_** + cos *n*θ_i_**; β_r*j*_ and γ_r*j*_ are the estimated regression coefficients for the *r*^th^ sin and cosine term respectively; and ε_ij_ is the error term.

With a single pair of Fourier terms (*n*=1), this gives a sinusoidal pattern of variation, with a single maximum and minimum whose phase (position in the year) and amplitude (distance between methylation maximum and minimum) is determined by β_1_ and γ_1,_ with the constraint that the maximum and minimum are 6 months apart. More complex patterns of seasonal variation are afforded by higher frequency pairs of Fourier terms (*r*>1).

For this analysis we modelled the effect of date of conception using a single pair of Fourier terms (*n*=1) and assessed goodness of fit by comparing full and covariate-only models using likelihood ratio tests. For both cohorts, covariates included child sex, and the first six principal components (PCs) obtained from unsupervised principal component analysis (PCA) of the normalised methylation M-values. The PCs were used to account for unmeasured and measured technical variation (due to bisulfite conversion sample plate, array slide etc) and cell composition effects (see Supplementary Tables 16 and 17). Additional checks confirmed no seasonal variation in estimated white cell composition in either cohort (see below). 450k Sentrix Column was included as an additional adjustment covariate for the ENID cohort since this was not robustly captured by any of the first 6 PCs (Supplementary Table 16). Child age was included as an additional adjustment covariate for the EMPHASIS cohort, since child ages ranged from 8 to 9 years, as was maternal nutritional intervention group (see Chandak *et al*^10^ and Saffari *et al*^65^ for further details).

For each CpG *j*, coefficient estimates β_j_, γ_j_ were determined by fitting regression models using *lm()* in R. Model goodness-of-fit was determined by likelihood ratio test (LRT) using *lrtest*() in R, comparing the full model including Fourier terms, with a baseline covariates-only model. A model p-value, p_*j*_ was then derived from the corresponding LRT chi-squared statistic. Thus for a given threshold, α, p_j_< α supports rejection of the null hypothesis that for CpG *j*, the full model including the effect of seasonality modelled by one pair of Fourier terms, fits no better than the covariate-only model at the α level.

To reduce the influence of methylation outliers, methylation values greater than three times the methylation inter-quartile range (IQR) beyond the 25^th^ or 75^th^ centiles were excluded prior to fitting the models. This resulted in 382,095 (267,369) outliers, or an average of 0.98 (0.68) outliers per CpG being removed from the ENID and EMPHASIS methylation datasets respectively.

### Inflation of test statistics

The concept of genomic inflation rests on the assumption that a relatively small number of loci will be associated with the exposure (or disease/outcome) of interest. Test statistics for the ENID (2yr) cohort did show signs of genomic inflation (lambda = 1.33). A similar level of inflation has been observed before in a study looking at the effect of periconceptional folate on DNAm (Gonseth et al^50^; lambda = 1.30 in the high folate exposure ‘Set 1’ analysis). There is also evidence of global and/or multi-locus effects of folate in other studies, including an RCT of folate supplementation in preganancy^51^, and there are many other examples including studies investigating the effect of mutations in the *MTHFR* gene (see review by Crider et al^76^). Gonseth et al use other lines of evidence to argue that inflation is likely due to a global effect of periconceptional maternal folate on epigenome-wide DNAm. Lambda for the Gambian EMPHASIS (8-9yr) cohort analysis is also much lower at 1.06, as we would expect on the assumption that SoC effects decrease with age. While we do not know if the SoC associations we observe in our cohorts are driven by seasonal differences in folate, we do observe significant seasonal differences in multiple one-carbon (C1) metabolites including folate in our population^8,12^, so that SoC may serve as a proxy for multi-locus C1 metabolite effects in our analyses. It is also possible that SoC serves as a proxy for other seasonal exposures. Our analysis of random and matched controls along with evidence for enrichment of loci with evidence of establishment of methylation states at periconception, further supports the notion that SoC-associations are not driven by population stratification or other spurious effects. We therefore did not consider it appropriate to adjust for ‘inflation’ of SoC-association test statistics.

### Identification of SoC-CpGs

For the ENID cohort, p-values, p_j_, were used to compute a false discovery rate for each CpG accounting for multiple testing (assuming 391,814 independent tests corresponding to the number of loci in array background) using *p.adjust()* in R with method= ‘fdr’. 768 CpGs with a FDR<5% formed the set of ‘**SoC-associated CpGs’**. Following the rationale described in the main text, SoC-associated CpGs with a SoC amplitude < 4% in the ENID cohort were then excluded to form the final set of 259 ‘**SoC-CpGs**’ (see Table 2).

### Selection of control CpGs

SoC-CpGs are enriched for intermediate methylation states (Fig. 3A), so that there is a risk that some downstream analyses reflect the distributional properties of these loci, rather than factors associated with their putative establishment at periconception. For this reason we identified a set of ‘**matched control**’ CpGs that were selected to have similar methylation Beta distributions to SoC-CpGs. Matched controls were drawn from array background (excluding SoC-CpGs and known MEs/ESS/SIV CpGs), with one matched control identified for each of the 768 SoC-associated CpGs (and 259 SoC-CpGs). Alignment of control and SoC-CpG methylation distributions was achieved using a two-sided Kolmogorov-Smirnov test for divergence of cumulative distribution functions (*ks.test()* in R) with a p-value threshold p>0.1. Examples are given in Supplementary Fig. 15, along with a comparison of sample mean distributions.

An additional 259 **random control** CpGs were randomly sampled from array background, again excluding SoC-CpGs and known MEs/ESS/SIV CpGs.

### CpG sets considered in analyses

Summary information on external datasets considered in the analyses is provided in Table 3. Further information on these is provided below.

i. *1,881 putative ME* CpGs overlapping array background from one or more of the following curated sets of loci: putative MEs exhibiting SIV identified in a multi-tissue WGBS screen in individuals of European and African-American decent described in Kessler *et al*^14^; and CpGs exhibiting ‘epigenetic supersimilarity’ and/or SIV derived from analysis of 450k data from individuals of European decent, as described in Van Baak *et al*^15^.
ii. *699 parent-of-origin-specific CpGs (PofOm CpGs)* overlapping array background from 229 regions with PofOm identified in Supplementary Table 1 from Zink et al^23^. PofOm identified in peripheral blood from Icelandic individuals.
iii. *RRBS early stage embryo data* from Chinese embryos described in Guo *et al*^4^ downloaded from GEO (accession number GSE49828). Only CpGs covered at ≥ 10x in pre-gastrulation inner cell mass and/or post-gastrulation embryonic liver were considered in this analysis. Further details are provided in Kessler *et al*^14^.
iv. *Sperm methylation data* from Japanese donors described in Okae *et al*^20^ downloaded from the Japanese Genotype-phenotype Archive (accession number S00000000006). Only CpGs covered at ≥ 10x were considered in this analysis.
v. *Germline gDMRs* (gDMRs), defined as contiguous 25 CpG regions that were hypomethylated (DNAm mean +1SD < 25%) in one gamete and hypermethylated (DNAm mean-1SD > 75%) in the other, were previously identified from WGBS data by Sanchez-Delgado *et al*^24^. Persistence of PofOm to the blastocyst and placental stages was established by identifying overlapping intermediately methylated regions in the relevant embryonic tissues, with confirmation of PofOm expression at multiple DMRs^24^. Japanese and US donors. See Sanchez-Delgado *et al*^24^ for further details.
vi. *Transposable elements (ERVs)* determined by RepeatMasker were downloaded from the UCSC hg19 annotations repository.
vii. ZFP57, TRIM28 and CTCF transcription factor binding sites identified from ChIP-seq in human embryonic kidney and human embryonic stem cells used in this analysis are described in Kessler *et al*^14^.

### Cluster-based adjustments

Many SoC-CpGs cluster together and this could influence some analyses. For example, methylation at CpGs may be highly correlated, which could influence comparisons of inter-CpG correlations between SoC-CpGs and controls (Fig. 3D). Also enrichment tests are likely to be influenced by neighbouring CpGs that together constitute a single ‘enrichment signal’ proximal to a particular genomic feature (e.g. a transcription factor binding site).

To account for this, cluster-adjusted analyses used ‘de-clustered’ CpG sets that were constructed as follows:

i. Create CpG clusters formed from adjacent CpGs where each CpG is within 5kbp of the nearest neighbouring CpG (see Supplementary Fig. 3 for a justification of this threshold);
ii. Construct de-clustered test set by randomly sampling a single CpG from each cluster; non-clustered ‘singleton’ CpGs are always selected.

In the case of SoC-CpGs, the set of 259 non-clustered CpGs were reduced to 161 CpGs after de-clustering.

### Chromatin state and histone (H3K) mark analysis

Data on chromatin states predicted by the ChromHMM 15-state model^77^ in H1 ESCs, and 3 fetal tissues (brain, muscle and small intestine) derived from 3 different germ layers, generated by the Roadmap Epigenomics Consortium^25^ was downloaded from the Washington University Roadmap Epigenomics repository. 15-state model predictions were collapsed to 8 states for visualisation purposes, with sub-classifications described in the Fig. 5 caption.

Overlaps with H3K marks for each of the above tissues were assessed using the *annotatr (v1.10.0)* package in R to interrogate the same Roadmap Epigenomics ChIP-seq data used by ChromHMM for chromatin state prediction.

### Additional modelling of seasonal variation in blood cell composition

Cell count estimates using the Houseman method^78^ were obtained using the *estimateCellCounts*() from *minfi* (v1.30.0) in R. Seasonal variation in blood cell composition was then modelled by Fourier regression with one pair of Fourier terms and sex (ENID+EMPHASIS) and age (EMPHASIS only) as adjustment covariates. Fitted models indicated no marked seasonal differences within and between cohorts (Supplementary Fig. 16).

### Genetic association analyses

#### Genotype data

mQTL and related SoC-association analyses were performed on all 284 individuals from the EMPHASIS (8-9yr) cohort for which we had QC’d genotype data. 259 SoC-CpGs, plus sets of 259 matched and random control CpGs were considered in this analysis. Subjects were genotyped using the Illumina Infinium Global Screening Array-24 v1.0 Beadchip (Illumina, California, U.S.) following standard protocols. Array-derived genotypes were pre-phased using SHAPEITv2 and imputation was performed using IMPUTEv2.3.2 on 1000 genomes phase 3 data. Further details are provided in Saffari et al^65^. SNPs with a MAF ≤ 10% were excluded, along with those with an IMPUTE ‘info’ metric ≤ 0.9, a stringent threshold to ensure maximum confidence in imputation quality. Imputed SNPs were then pruned (using plink v1.90 -indep-pairwise with window size 50, step size 5 and r^2^ threshold of 0.8) to remove SNPs in strong LD. Finally, to minimise the influence of low frequency homozygous variants in linear models, analysis was restricted to SNPs with 10 or more homozygous variants, resulting in a final dataset comprising 2,609,310 SNPs.

#### Identification of mQTLs and ‘GxE’ SNPs

mQTL analysis was performed using the *GEM* package (v1.10.0) from R Bioconductor^79^. SNP effects on methylation were modelled as follows:

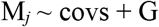

Where M_*j*_ is the methylation M-value for CpG *j*, G is the SNP genotype coded as allelic dosage (0,1,2) and covs correspond to the adjustment covariates used in the main EMPHASIS analysis (PCs 1-6, child age, sex and intervention status).

GxE (SoC) effects were modelled as follows:

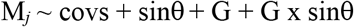

when sinθ is the most significant Fourier term in the main SoC analysis,
OR

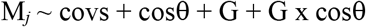

when cosθ is the most significant Fourier term in the main SoC analysis.

Here, G and covs are as described above.

Since most mQTL effects are known to act in *cis*^28^, in order to maximise power a two-step procedure was used to identify significant mQTLs:

i. *Identification of cis-mQTLs* passing FDR<5% considering the reduced set of SNPs within 1Mbp of a CpG in the set to be analysed (SoC-CpGs, matched or random controls);
ii. *Identification of trans-mQTLs* passing FDR<5%, considering the full set of 2.6M SNPs.

#### Calculation of methylation variance explained by mQTLs

For each CpG *j*, total methylation variance explained was calculated for the following model:

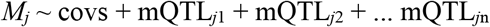

where covs are as defined above, and mQTL_*jn*_ is the genotype (coded 0,1,2) of the *n*th mQTL mapping to CpG *j.* Methylation variance explained was calculated from the model adjusted R^2^ value, adjR^2^_mQTL_, to account for different levels of model complexity due to the differing number of mQTL identified for each CpG.

A final estimate of mQTL methylation variance explained was obtained by subtracting variance explained by the covariate-only model:

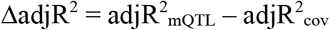

where adjR^2^ is the adjusted R^2^ for the covariate-only model,

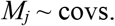

#### SoC association analysis

Potential confounding of SoC-DNAm signals by SoC-associated genetic variants was assessed by analysing SoC associations with all 2,771 SoC-CpG-associated mQTLs identified in the mQTL analysis (see Table 4). SoC association analysis was performed under an allelic model using -assoc with *plink* v1.90. Significant mQTL-SoC associations were identified using FDR<5% which assumes independence of all mQTL-SoC associations. We also considered a more liberal multiple testing correction threshold: *p_Bonf_* = 0.05/130, assuming complete dependence of all *cis-*mQTL mapping to each of the 130 SoC-CpG with an associated mQTL.

#### Genetic ancestry sensitivity analysis

An investigation of population structure in the EMPHASIS cohort was conducted by first performing a principal component analysis (PCA) using -pca in plink v1.90. The PCA was performed on non-imputed data, with LD pruning using -indep-pairwise 50 5 0.2 in plink v1.90, corresponding to an r^2^ threshold of 0.2^80^. Evidence for population structure was then obtained by plotting the first four principal components (Supplementary Fig. 12A left). Confirmation of a likely link to Gambian ethnic ancestry in this largely ethnically Mandinka region of Gambia followed our observation that a distinct cluster (Supplementary Fig. 12B) was primarily made up of individuals from a single, predominantly ethnic Fula village.

A sensitivity analysis to check the effect of accounting for ancestry differences was performed by repeating the main analysis with the following additional covariates:

1. EMPHASIS (8-9yr) cohort

We adjusted for genetic ancestry directly using the first two genetic principal components identified in the genetic PCA.

2. ENID (2yr) cohort

Since no genetic data is available for this cohort, and since individuals from this cohort are drawn from the same villages as the EMPHASIS cohort on which we did the genetic PCA, we reasoned that village of origin is a useful proxy for genetic ancestry in our population. We therefore included an additional covariate dichotomised according to whether or not an individual was one of the 9 who came from the predominantly Fula village identified as genetic outlier in the EMPHASIS PCA analysis.

#### Sensitivity analysis to investigate confounding by infant sex

Our finding that multiple SoC-CpGs were associated with infant sex in previous EWAS prompted us to perform a sensitivity analysis checking for the possibility of a residual confounding effect due to sex. To do this we regressed out the effect of infant sex at each CpG in the M-value methylation matrix, prior to the main regression analysis. We then re-ran the Fourier regression analysis with and without an additional adjustment for infant sex in Fourier regression models. As expected, given that we adjusted for infant sex in the main analysis, this produced near identical results, suggesting that the main analysis was not confounded by infant sex.

### All bootstrapped confidence intervals presented in this paper use 1,000 bootstrap samples

#### Ethics approval and consent to participate

Ethics approval for the Gambian ENID and EMPHASIS trials was obtained from the joint Gambia Government/MRC Unit The Gambia’s Ethics Committee (ENID: SCC1126v2; EMPHASIS: SCC1441). The ENID study is registered as ISRCTN49285450. The EMPHASIS study is registered as ISRCTN14266771. Signed informed consent for both studies was obtained from parents, and verbal assent was additionally obtained from the older children who participated in the EMPHASIS study.

## Funding

The Gambian ENID trial was jointly funded by the UK Medical Research Council (MRC) and the Department for International Development (DFID) under the MRC/DFID Concordat agreement (MRC Program MC-A760-5QX00). Methylation analysis of ENID samples was supported by the Bill & Melinda Gates Foundation (grant no: OPP1 066947). The Gambian EMPHASIS study is jointly funded by MRC, DFID and the Department of Biotechnology, Ministry of Science and Technology, India under the Newton Fund initiative (MRC grant no.: MR/N006208/1 and DBT grant no.: BT/IN/DBT-MRC/DFID/24/GRC/2015–16). Further support for this analysis was provided by MRC Grant MR/M01424X/1. We acknowledge the work of the full EMPHASIS Study Group (www.emphasisstudy.org) in acquiring this data.

## Author contributions

MJS conceived the study, performed the bulk of the analysis and wrote the original draft. AS, NJK, MD and PTJ performed additional analyses. PI, AD and MB contributed to sample acquisition and/or processing. MJS and AMP obtained the funding. GRC, CHDF, MJS, SEM, MR, ZH, CC, DM and AMP established the cohorts analysed in this study and/or provided data used in the analysis. All authors contributed to and approved the final manuscript.

## Disclaimer

Where authors are identified as personnel of the International Agency for Research on Cancer / World Health Organization, the authors alone are responsible for the views expressed in this article and they do not necessarily represent the decisions, policy or views of the International Agency for Research on Cancer / World Health Organization.

## Competing interests

Authors declare that they have no competing interests.

## Data and materials availability

### Data and code availability

ENID 450k methylation data analysed for this study is deposited in GEO (GSE99863). Gambian EMPHASIS EPIC methylation data is available on request and will be made publicly available once results from the main EMPHASIS study have been published. Sources and locations of other publicly available data used in this analysis are described in METHODS. Bespoke code used in the analysis is available at https://github.com/mattjsilver/SoCFourier.

## Notes

### Competing Interest Statement

The authors have declared no competing interest.

### Summary of Updates

1. Prompted by evidence of a decline in SoC effect size with age, we have applied a more stringent threshold for identifying SoC-CpGs (FDR<5%), and have shifted from a discovery-replication study design to one in which we use triangulation of evidence between cohorts to strengthen evidence for consistency of SoC effects, while highlighting SoC hotpots at age 2 as robust sentinels of periconceptional exposure. 2. We have removed the eForge histone mark analysis using in the light of poor correspondence between eForge H3K enrichments and H3K genome browser peaks and replaced this with a chromatin state analysis and direct interrogation of Epigenome Roadmap H3K marks in H1 ES cells and fetal tissues from each germ layer. 3. We have changed the way we analyse proximal ERVs and transcription factor binding sites to include a wider range of proximity distances. 4. We have increased the power of the mQTL analysis by performing separate investigations of mQTL effects in cis and in trans. 5. We have changed our strategy for analysing GxE effects after noting that our previous method had poor control of false positives. 6. We have assessed the potential influence of genetic ancestry. 6. We have compared our findings with those from related studies and with previous results from EWAS and GWAS.

## References

1. Smith, Z. D. & Meissner, A. DNA methylation: roles in mammalian development. Nature Reviews Genetics 14, 204–220 (2013).

2. Jeltsch, A. Molecular Enzymology of Mammalian DNA Methyltransferases. in DNA Methylation: Basic Mechanisms 203–225 (Springer-Verlag). doi:10.1007/3-540-31390-7_7.

3. Feldmann, A. et al. Transcription Factor Occupancy Can Mediate Active Turnover of DNA Methylation at Regulatory Regions. PLoS Genetics 9, (2013).

4. Guo, H. et al. The DNA methylation landscape of human early embryos. Nature 511, 606–610 (2014).

5. Fleming, T. P. et al. Origins of lifetime health around the time of conception: causes and consequences. The Lancet 391, 1842–1852 (2018).

6. Birney, E., Smith, G. D. & Greally, J. M. Epigenome-wide Association Studies and the Interpretation of Disease -Omics. PLOS Genetics 12, e1006105 (2016).

7. Moore, S. E. et al. Prenatal or early postnatal events predict infectious deaths in young adulthood in rural Africa. International journal of epidemiology 28, 1088–95 (1999).

8. Dominguez-Salas, P. et al. DNA methylation potential: Dietary intake and blood concentrations of one-carbon metabolites and cofactors in rural African women. American Journal of Clinical Nutrition 97, 1217–1227 (2013).

9. Moore, S. E. et al. A randomized trial to investigate the effects of pre-natal and infant nutritional supplementation on infant immune development in rural Gambia: the ENID trial: Early Nutrition and Immune Development. BMC pregnancy and childbirth 12, 107 (2012).

10. Chandak, G. R. et al. Protocol for the EMPHASIS study; epigenetic mechanisms linking maternal pre-conceptional nutrition and children’s health in India and Sub-Saharan Africa. BMC Nutrition 3, 81 (2017).

11. Waterland, R. A. et al. Season of conception in rural gambia affects DNA methylation at putative human metastable epialleles. PLoS genetics 6, e1001252 (2010).

12. Dominguez-Salas, P. et al. Maternal nutrition at conception modulates DNA methylation of human metastable epialleles. Nature Communications 5, 1–7 (2014).

13. Silver, M. et al. Independent genomewide screens identify the tumor suppressor VTRNA2-1 as a human epiallele responsive to periconceptional environment. Genome Biology 16, 118 (2015).

14. Kessler, N. J., Waterland, R. A., Prentice, A. M. & Silver, M. J. Establishment of environmentally sensitive DNA methylation states in the very early human embryo. Science Advances 4, eaat2624 (2018).

15. Van Baak, T. E. et al. Epigenetic supersimilarity of monozygotic twin pairs. Genome Biology 19, 2 (2018).

16. Rakyan, V. K., Blewitt, M. E., Druker, R., Preis, J. I. & Whitelaw, E. Metastable epialleles in mammals. Trends in genetics: TIG 18, 348–51 (2002).

17. Anderson, O. S., Sant, K. E. & Dolinoy, D. C. Nutrition and epigenetics: an interplay of dietary methyl donors, one-carbon metabolism and DNA methylation. The Journal of Nutritional Biochemistry 23, 853–859 (2012).

18. Gunasekara, C. J. & Waterland, R. A. A new era for epigenetic epidemiology. Epigenomics 11, 1647–1649 (2019).

19. Rayco-Solon, P., Fulford, A. & Prentice AM. Differential effects of seasonality on preterm birth and intrauterine growth. American Journal of Clinical Nutrition 81, 134–139 (2005).

20. Okae, H. et al. Genome-Wide Analysis of DNA Methylation Dynamics during Early Human Development. PLoS Genetics 10, e1004868 (2014).

21. Monk, D., Mackay, D. J. G., Eggermann, T., Maher, E. R. & Riccio, A. Genomic imprinting disorders: lessons on how genome, epigenome and environment interact. Nature Reviews Genetics 20, 235–248 (2019).

22. James, P. et al. Candidate genes linking maternal nutrient exposure to offspring health via DNA methylation: a review of existing evidence in humans with specific focus on one-carbon metabolism. International Journal of Epidemiology 1–28 (2018) doi:10.1093/ije/dyy153.

23. Zink, F. et al. Insights into imprinting from parent-of-origin phased methylomes and transcriptomes. Nature Genetics 50, 1542–1552 (2018).

24. Sanchez-Delgado, M. et al. Human Oocyte-Derived Methylation Differences Persist in the Placenta Revealing Widespread Transient Imprinting. PLOS Genetics 12, e1006427 (2016).

25. Kundaje, A. et al. Integrative analysis of 111 reference human epigenomes. Nature 518, 317–330 (2015).

26. Waterland, R. A. & Jirtle, R. L. Transposable elements: targets for early nutritional effects on epigenetic gene regulation. Molecular and cellular biology 23, 5293–300 (2003).

27. Kazachenka, A. et al. Identification, Characterization, and Heritability of Murine Metastable Epialleles: Implications for Non-genetic Inheritance. Cell 1–13 (2018) doi:10.1016/j.cell.2018.09.043.

28. Gaunt, T. R. et al. Systematic identification of genetic influences on methylation across the human life course. Genome Biology 17, 61 (2016).

29. Hennig, B. J. et al. Cohort Profile: The Kiang West Longitudinal Population Study (KWLPS)— a platform for integrated research and health care provision in rural Gambia. International Journal of Epidemiology dyv206 (2015) doi:10.1093/ije/dyv206.

30. Ba, Y. et al. Relationship of folate, vitamin B12 and methylation of insulin-like growth factor-II in maternal and cord blood. Eur J Clin Nutr 65, 480–485 (2011).

31. Tobi, E. W. et al. Prenatal Famine and Genetic Variation Are Independently and Additively Associated with DNA Methylation at Regulatory Loci within IGF2/H19. PLoS One 7, (2012).

32. Buniello, A. et al. The NHGRI-EBI GWAS Catalog of published genome-wide association studies, targeted arrays and summary statistics 2019. Nucleic Acids Res 47, D1005–D1012 (2019).

33. Do, C. et al. Genetic-epigenetic interactions in cis: A major focus in the post-GWAS era. Genome Biology 18, 1–22 (2017).

34. Comuzzie, A. G. et al. Novel Genetic Loci Identified for the Pathophysiology of Childhood Obesity in the Hispanic Population. PLOS ONE 7, e51954 (2012).

35. Evans, D. S. et al. Fine-mapping, novel loci identification, and SNP association transferability in a genome-wide association study of QRS duration in African Americans. Hum Mol Genet 25, 4350–4368 (2016).

36. van der Harst, P. et al. 52 Genetic Loci Influencing Myocardial Mass. J Am Coll Cardiol 68, 1435–1448 (2016).

37. Kühnen, P. et al. Interindividual Variation in DNA Methylation at a Putative POMC Metastable Epiallele Is Associated with Obesity. Cell metabolism 24, 502–509 (2016).

38. Molaro, A. et al. Sperm methylation profiles reveal features of epigenetic inheritance and evolution in primates. Cell 146, 1029–1041 (2011).

39. Hammoud, S. S. et al. Distinctive chromatin in human sperm packages genes for embryo development. Nature 460, 473–478 (2009).

40. Rada-Iglesias, A. et al. A unique chromatin signature uncovers early developmental enhancers in humans. Nature 470, 279–285 (2011).

41. Vukic, M., Wu, H. & Daxinger, L. Making headway towards understanding how epigenetic mechanisms contribute to early-life effects. Philosophical Transactions of the Royal Society B: Biological Sciences 374, 20180126 (2019).

42. Simpkin, A. J. et al. Longitudinal analysis of DNA methylation associated with birth weight and gestational age. Human Molecular Genetics 24, 3752–3763 (2015).

43. Carpenter, B. L. et al. Mother–child transmission of epigenetic information by tunable polymorphic imprinting. Proceedings of the National Academy of Sciences 201815005 (2018) doi:10.1073/pnas.1815005115.

44. Bogutz, A. B. et al. Evolution of imprinting via lineage-specific insertion of retroviral promoters. Nature Communications 10, 1–14 (2019).

45. Cavalli, G. & Heard, E. Advances in epigenetics link genetics to the environment and disease. Nature 571, 489–499 (2019).

46. Imbeault, M., Helleboid, P. Y. & Trono, D. KRAB zinc-finger proteins contribute to the evolution of gene regulatory networks. Nature 543, 550–554 (2017).

47. Shi, H. et al. ZFP57 regulation of transposable elements and gene expression within and beyond imprinted domains. Epigenetics & Chromatin 12, 49 (2019).

48. Monteagudo-Sánchez, A. et al. The role of ZFP57 and additional KRAB-zinc finger proteins in the maintenance of human imprinted methylation and multi-locus imprinting disturbances. Nucleic Acids Res doi:10.1093/nar/gkaa837.

49. Amarasekera, M. et al. Genome-wide DNA methylation profiling identifies a folate-sensitive region of differential methylation upstream of ZFP57-imprinting regulator in humans. FASEB journal: official publication of the Federation of American Societies for Experimental Biology 1–9 (2014) doi:10.1096/fj.13-249029.

50. Gonseth, S. et al. Periconceptional folate consumption is associated with neonatal DNA methylation modifications in neural crest regulatory and cancer development genes. Epigenetics 10, 1166–76 (2015).

51. Irwin, R. E. et al. A randomized controlled trial of folic acid intervention in pregnancy highlights a putative methylation-regulated control element at ZFP57. Clinical Epigenetics 11, 31 (2019).

52. Finer, S. et al. Is famine exposure during developmental life in rural Bangladesh associated with a metabolic and epigenetic signature in young adulthood? A historical cohort study. BMJ Open 6, e011768 (2016).

53. Elliott, G. et al. Intermediate DNA methylation is a conserved signature of genome regulation. Nature Communications 6, 6363 (2015).

54. Tobi, E. W. et al. Selective Survival of Embryos Can Explain DNA Methylation Signatures of Adverse Prenatal Environments. Cell Reports 25, 2660–2667.e4 (2018).

55. Kang, L. et al. Aberrant allele-switch imprinting of a novel IGF1R intragenic antisense non-coding RNA in breast cancers. European Journal of Cancer 51, 260–270 (2015).

56. Sun, J. et al. A novel antisense long noncoding RNA within the IGF1Rgene locus is imprinted in hematopoietic malignancies. Nucleic Acids Research 42, 9588–9601 (2014).

57. Boucher, J. et al. Insulin and insulin-like growth factor 1 receptors are required for normal expression of imprinted genes. Proceedings of the National Academy of Sciences 111, 14512–14517 (2014).

58. Randhawa, R. & Cohen, P. The role of the insulin-like growth factor system in prenatal growth. Molecular Genetics and Metabolism 86, 84–90 (2005).

59. Aguirre, G. A., Ita, J. R., Garza, R. G. & Castilla-Cortazar, I. Insulin-like growth factor-1 deficiency and metabolic syndrome. Journal of Translational Medicine 14, 1–23 (2016).

60. Larsson, O., Girnita, A. & Girnita, L. Role of insulin-like growth factor I receptor signalling in cancer. British Journal of Cancer 92, 2097–2101 (2005).

61. Tsai, P.-C. et al. DNA Methylation Changes in the IGF1R Gene in Birth Weight Discordant Adult Monozygotic Twins. Twin Research and Human Genetics 18, 635–646 (2015).

62. Singmann, P. et al. Characterization of whole-genome autosomal differences of DNA methylation between men and women. Epigenetics & Chromatin 8, 43 (2015).

63. Shah, S. et al. Genetic and environmental exposures constrain epigenetic drift over the human life course. Genome Research 24, 1725–1733 (2014).

64. Suderman, M. et al. Sex-associated autosomal DNA methylation differences are wide-spread and stable throughout childhood. http://biorxiv.org/lookup/doi/10.1101/118265 (2017) doi:10.1101/118265.

65. Saffari, A. et al. Effect of maternal preconceptional and pregnancy micronutrient interventions on children’s DNA methylation: Findings from the EMPHASIS study. The American Journal of Clinical Nutrition nqaa193 (2020) doi:10.1093/ajcn/nqaa193.

66. Kuehnen, P. et al. An Alu Element–Associated Hypermethylation Variant of the POMC Gene Is Associated with Childhood Obesity. PLoS Genetics 8, e1002543 (2012).

67. James, P. T. et al. Maternal One-Carbon Metabolism and Infant DNA Methylation between Contrasting Seasonal Environments: A Case Study from The Gambia. Current Developments in Nutrition 3, nzy082 (2019).

68. Hernandez-Vargas, H. et al. Exposure to aflatoxin B 1 in utero is associated with DNA methylation in white blood cells of infants in The Gambia. International Journal of Epidemiology 44, 1238–1248 (2015).

69. Webster, A. P. et al. Increased DNA methylation variability in rheumatoid arthritis-discordant monozygotic twins. Genome Medicine 10, 1–12 (2018).

70. Ek, W. E. et al. Genetic variants influencing phenotypic variance heterogeneity. Human molecular genetics 27, 799–810 (2018).

71. Owens, S. et al. Periconceptional multiple-micronutrient supplementation and placental function in rural Gambian women: A double-blind, randomized, placebo-controlled trial. American Journal of Clinical Nutrition 102, 1450–1459 (2015).

72. Min, J. L., Hemani, G., Davey Smith, G., Relton, C. & Suderman, M. Meffil: efficient normalization and analysis of very large DNA methylation datasets. Bioinformatics (Oxford, England) 34, 3983–3989 (2018).

73. Chen, Y. et al. Discovery of cross-reactive probes and polymorphic CpGs in the Illumina Infinium HumanMethylation450 microarray. Epigenetics: official journal of the DNA Methylation Society 8, 203–9 (2013).

74. Fortin, J. P. et al. Functional normalization of 450k methylation array data improves replication in large cancer studies. Genome Biology 15, 0–42 (2014).

75. Du, P. et al. Comparison of Beta-value and M-value methods for quantifying methylation levels by microarray analysis. BMC bioinformatics 11, 587 (2010).

76. Crider, K. S., Yang, T. P., Berry, R. J. & Bailey, L. B. Folate and DNA Methylation: A Review of Molecular Mechanisms and the Evidence for Folate’s Role2. Adv Nutr 3, 21–38 (2012).

77. Ernst, J. & Kellis, M. ChromHMM: automating chromatin-state discovery and characterization. Nature Methods 9, 215–216 (2012).

78. Jaffe, A. E. & Irizarry, R. a. Accounting for cellular heterogeneity is critical in epigenome-wide association studies. Genome biology 15, R31 (2014).

79. Pan, H., Holbrook, J. D., Karnani, N. & Kwoh, C. K. Gene, Environment and Methylation (GEM): a tool suite to efficiently navigate large scale epigenome wide association studies and integrate genotype and interaction between genotype and environment. BMC Bioinformatics 17, 299 (2016).

80. Weale, M. E. Quality control for genome-wide association studies. Methods Mol. Biol. 628, 341–372 (2010).

