## Supplementary Figures for "Environmentally sensitive hotspots in the methylome of the early human embryo"

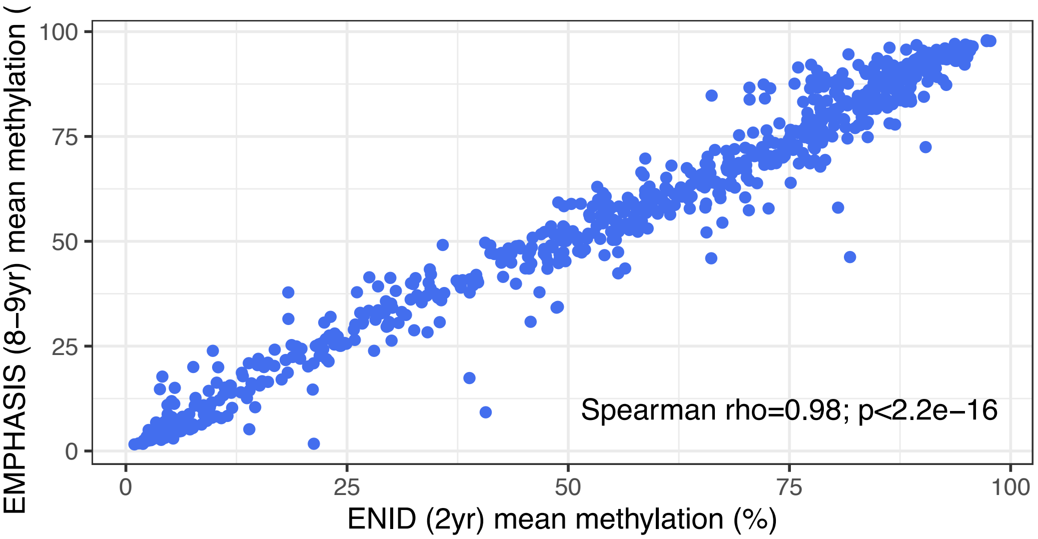


**Supplementary Figure 1. Cross-cohort correlation of mean methylation values at 768 SoC-associated loci.** Data comprises n=233 and n=289 individuals from the ENID and EMPHASIS cohorts respectively. 768 SoC-associated CpGs (FDR<5%) identified in the ENID cohort.

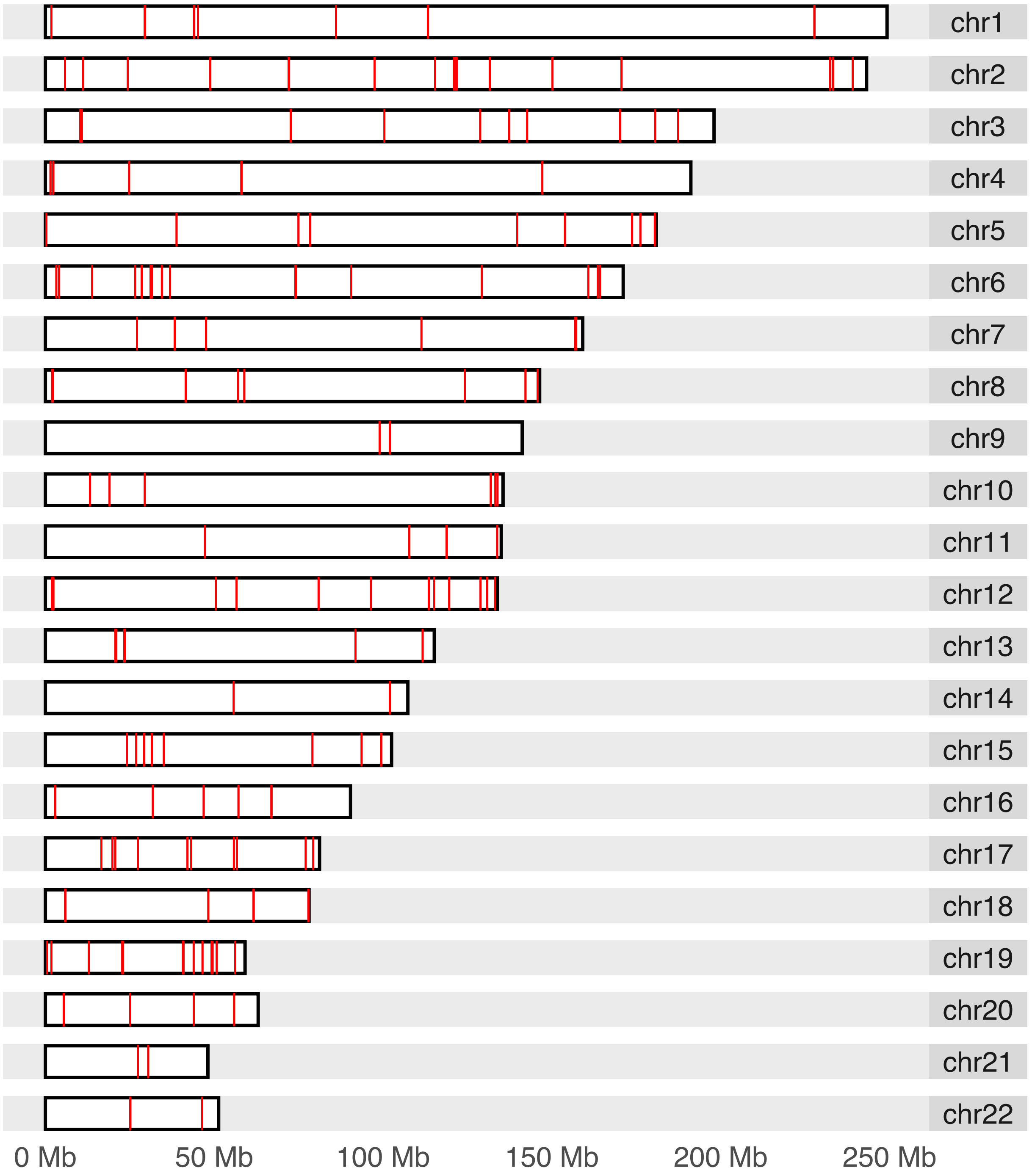


**Supplementary Figure 2. Genomic locations (hg19) of 259 SoC-CpGs.**
Data from Supplementary Table 4.

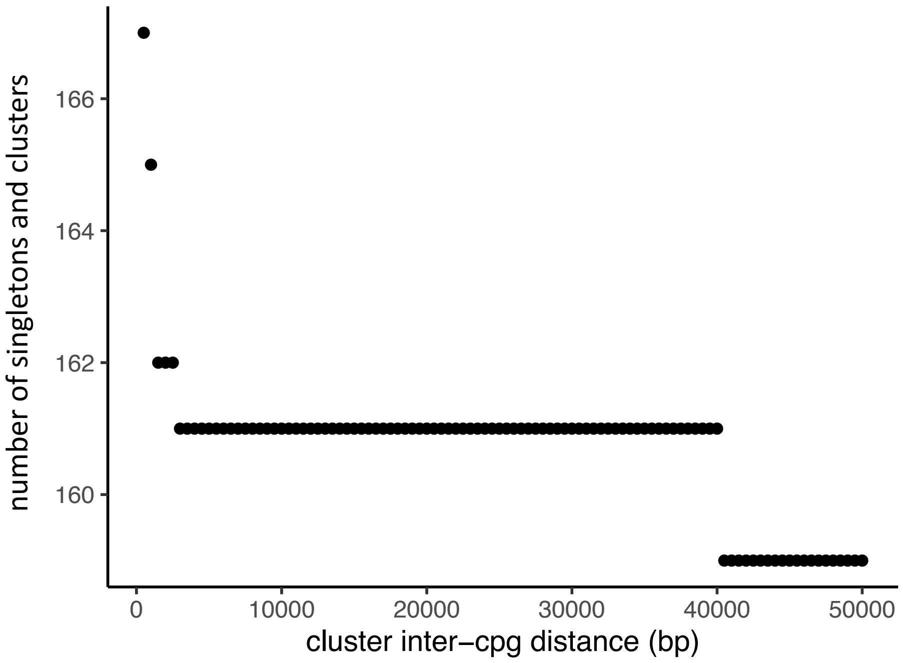


**Supplementary Figure 3.** Relationship between the inter-CpG distance used to define SoC-CpG clusters and the number of identified clusters and singletons (CpGs not falling within a cluster).

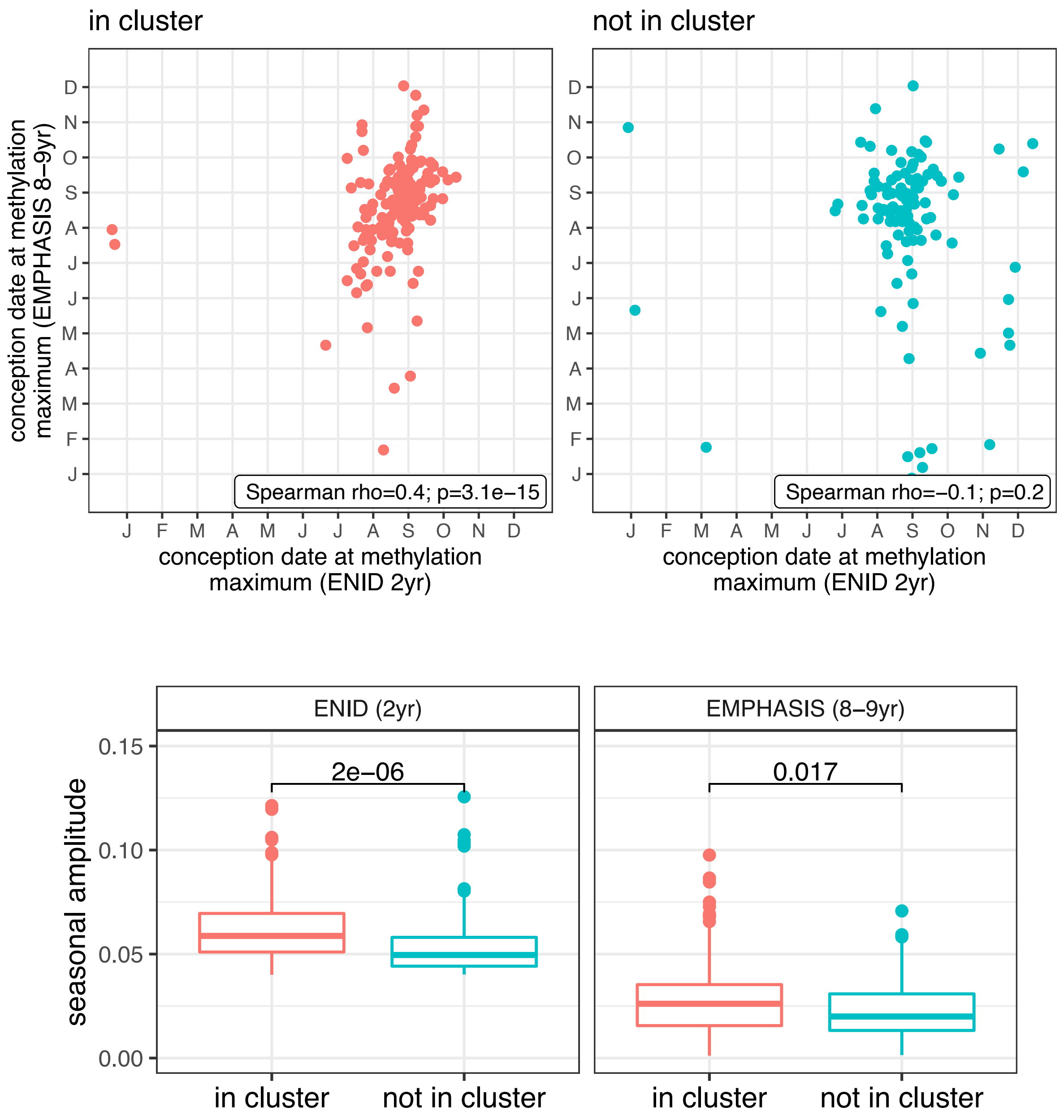


**Supplementary Figure 4. Comparison of SoC effects at SoC-CpGs within and outside of SoC-CpG clusters. Top:** Conception date at modelled methylation maxima for ENID (x-axis) and EMPHASIS (y-axis) cohorts, according to whether locus falls within (n=154; left-red), or outside (n=105; right-blue) of a SoC-CpG cluster. **Bottom:** Seasonal effect size (amplitude) for ENID (left) and EMPHASIS (cohorts), according to whether locus falls within (red), or outside (blue) of a SoC-CpG cluster. Numbers are P-values for two-sided Wilcoxon Rank Sum tests under the null hypothesis of no difference between the two distributions.

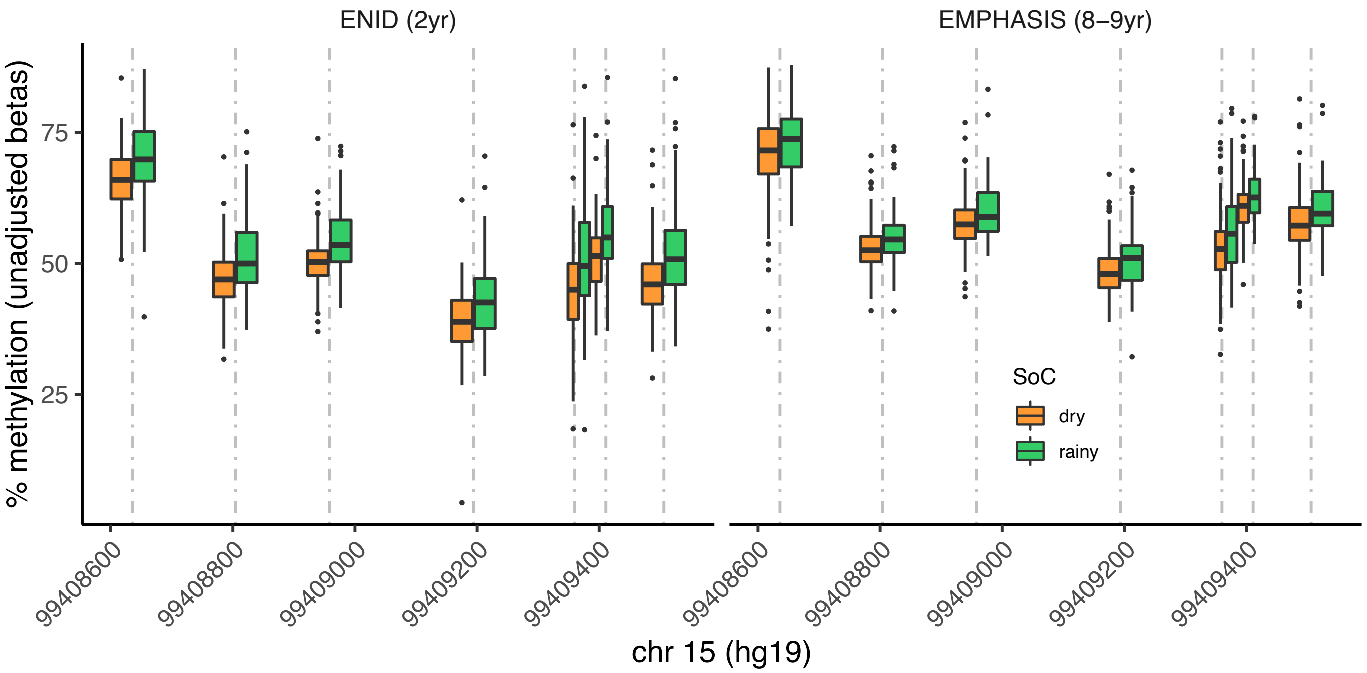


**Supplementary Figure 5. Seasonal variation at the *IGF1R* locus in ENID and EMPHASIS cohorts.** Unadjusted methylation Beta values are plotted. Boxes represent IQRs for season-specific DNAm at each locus. Note that for visualisation purposes Gambian seasons are dichotomised (Rainy: Jan-Jun; Dry: Jul-Dec), whereas seasonality is modelled as a continuous cyclical variable in Fourier regression models used in the main analysis.

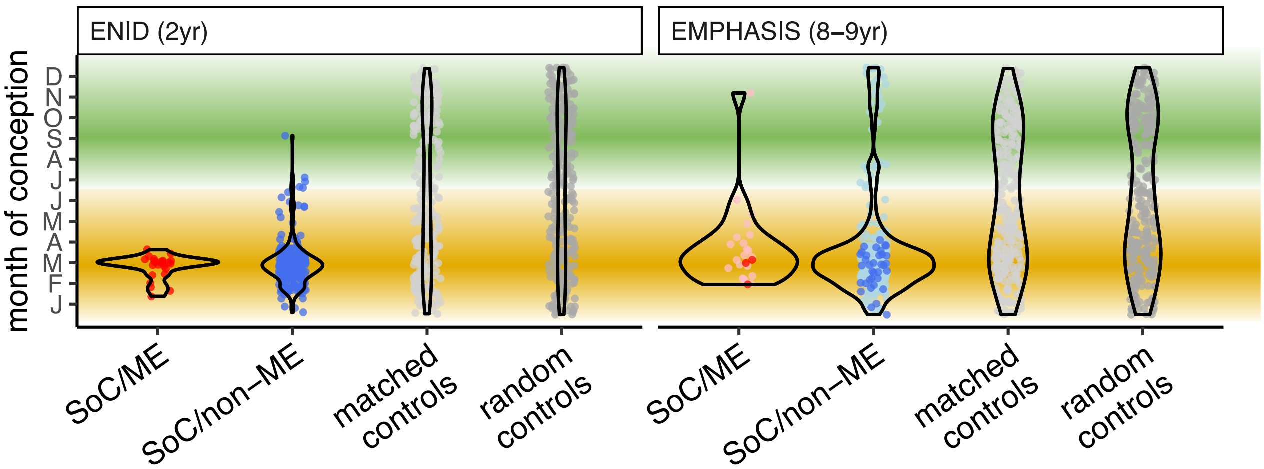


**Supplementary Figure 6. Date of conception at modelled methylation minima.** Conception dates at methylation minima for loci plotted in Fig. 3C (top). Note that since seasonality is modelled by a single pair of Fourier terms, maxima and minima are 6 months apart.

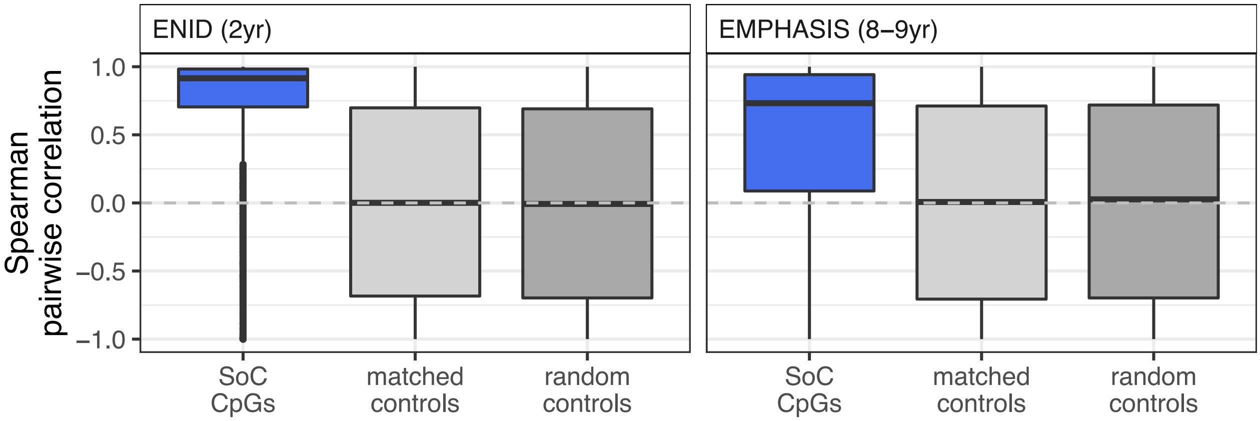


**Supplementary Figure 7. Cluster-adjusted pairwise inter-CpG methylation correlations for CpG sets in ENID (left) and EMPHASIS (right) cohorts.** This is the same as Fig. 3D, but with a single CpG randomly sampled from each CpG cluster so that CpGs in each set are a minimum distance of 5000bp apart. This shows that pairwise correlation distributions are not disproportionately driven by large numbers of pairwise correlations between highly-correlated neighbours.

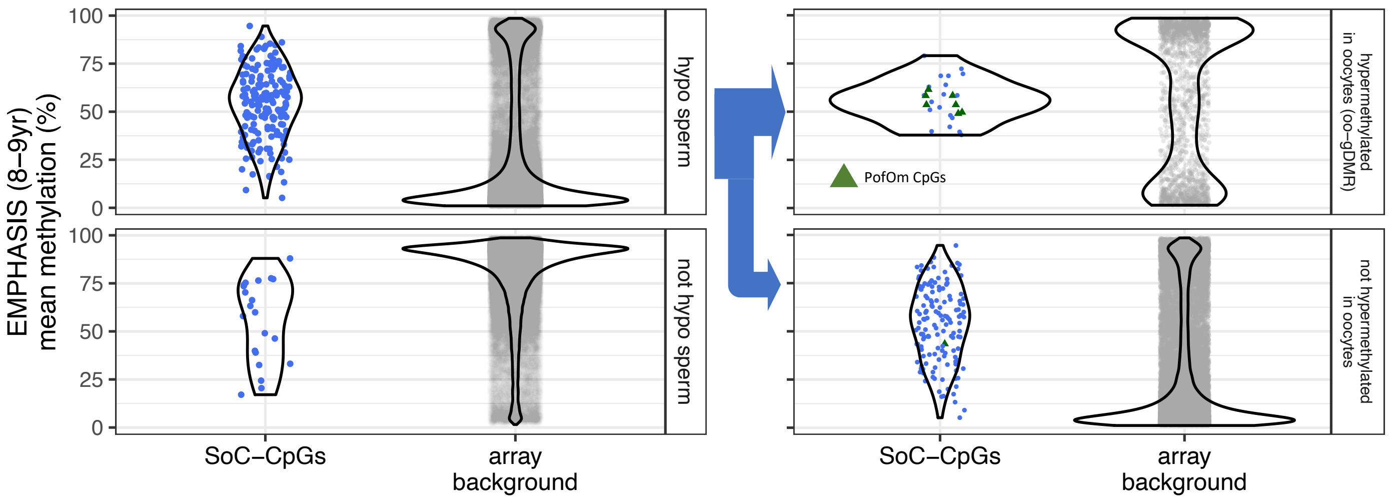


**Supplementary Figure 8. Mean methylation at SoC-CpGs and array background measured in n=289 individuals in the EMPHASIS (8-9yr) cohort stratified by sperm and oocyte methylation status**. (**Left**): Methylation stratified according to sperm methylation status reported in Okae *et al.*[20]. Sperm hypomethylation is defined as methylation ≤ 25%.(**Right**) as left but with loci hypomethylated in sperm further stratified according to oocyte gDMR (oo-gDMR) status reported in Sanchez-Delgado *et al*[24]. PofOm CpGs are those identified in Zink *et al.*[24] oo-gDMR have oocyte methylation >75%.

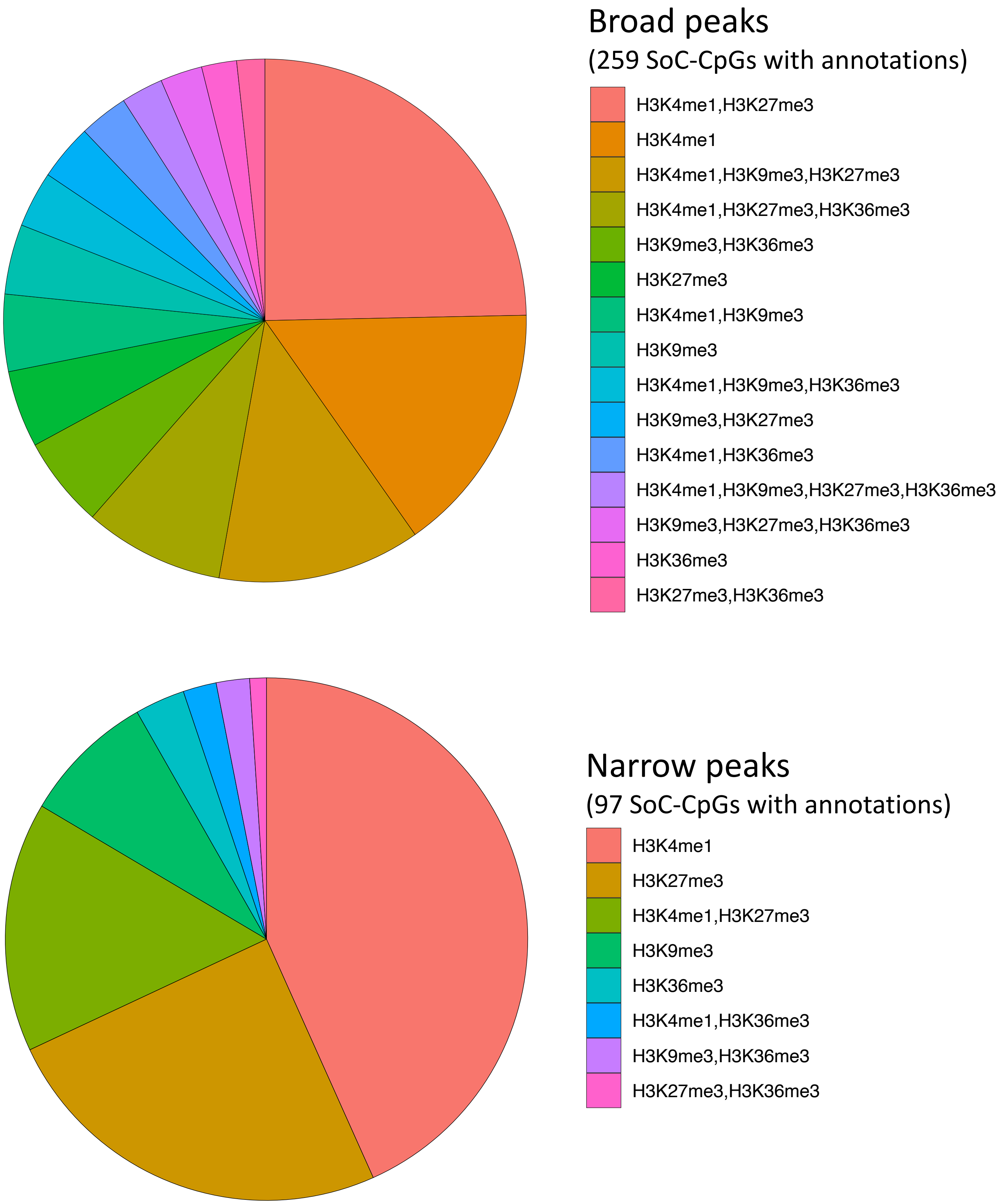


**Supplementary Figure 9. Histone marks overlapping SoC-CpGs in H1 embryonic stem cells**. Histone marks or combinations thereof are ordered by abundance. H3 marks were generated by the Roadmap Epigenomics Consortium[25] and downloaded using the *annotatr (v1.10.0)* package in R. Broad peaks overlapping all 259 SoC-CpGs (**top**) and (more sharply defined) narrow peaks overlapping 97 SoC-CpGs (**bottom**) correspond to different thresholds used by the MACS peak-calling algorithm. See Kundaje et al[25] for further details.

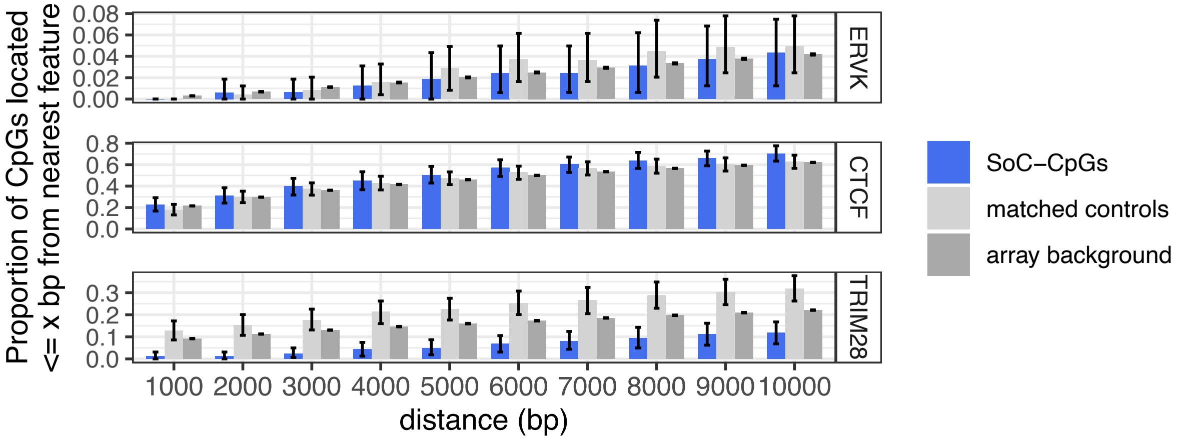


**Supplementary Figure 10. Links between ERVK endogenous retroviral elements, ZFP57 and TRIM28 binding sites, and DNAm at SoC-CpGs. (A)** Proportion of SoC-CpGs, matched controls and array background CpGs proximal to ERVK endogenous retroviral elements (top), and ZFP57 and TRIM28 binding sites (bottom), within the specified distance. CpG clustering effects are removed by sampling a single CpG from each cluster (see Methods). Error bars are bootstrapped 95% CIs.

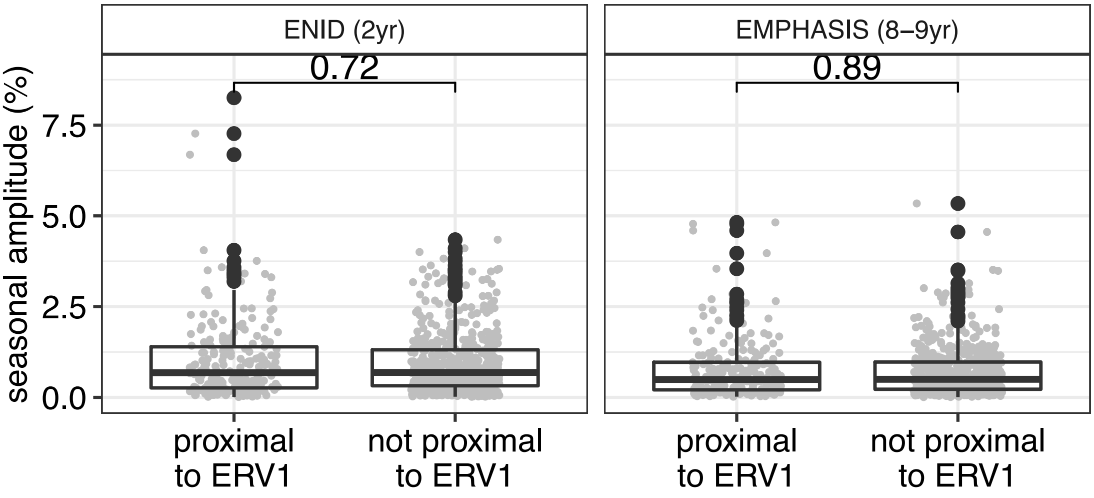


**Supplementary Figure 11.** Difference in SoC effect size (seasonal amplitude) between matched control loci proximal (<10kbp) to ERV1 (n=213) and not (n=555) in the ENID and EMPHASIS cohorts. P-values refer to two-sided Wilcoxon Rank Sum tests under the null hypothesis of no difference between the two distributions tested Controls match the DNAm distributions of n=768 SoC-associated loci in the ENID cohort (FDR<5%).

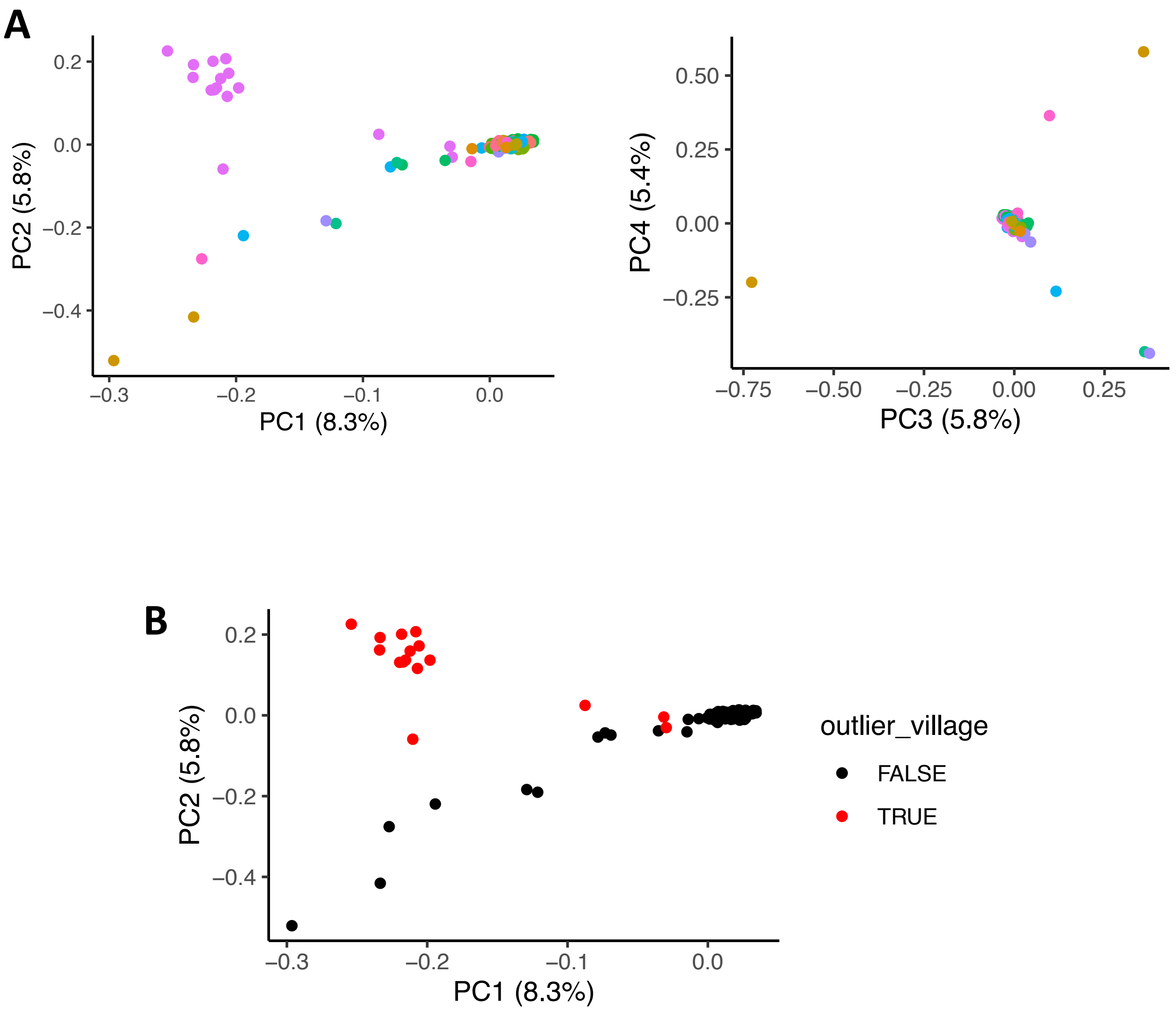


**Supplementary Figure 12. Population structure in the EMPHASIS cohort.**
**A:** 1st and 2nd (left) and 2nd and 3rd (right) principal components (PCs) from a PCA of genome-wide genetic data from n=294 individuals from the EMPHASIS cohort. Figures in brackets give % variance explained by each PC. Colours correspond to the 24 villages from the Kiang West region which are covered in this cohort. A majority of individuals are of Mandinka origin and villages that are predominantly Mandinka are tightly clustered in the PCA plot. The distinct cluster in the top left of the first plot corresponds to individuals from a single village which is predominantly Fula. **B:** Same as **A** left, but with individuals from the 'outlier village' plotted as a distinct colour. This illustrates that all 16 individuals from this village are distinct from the main cluster.

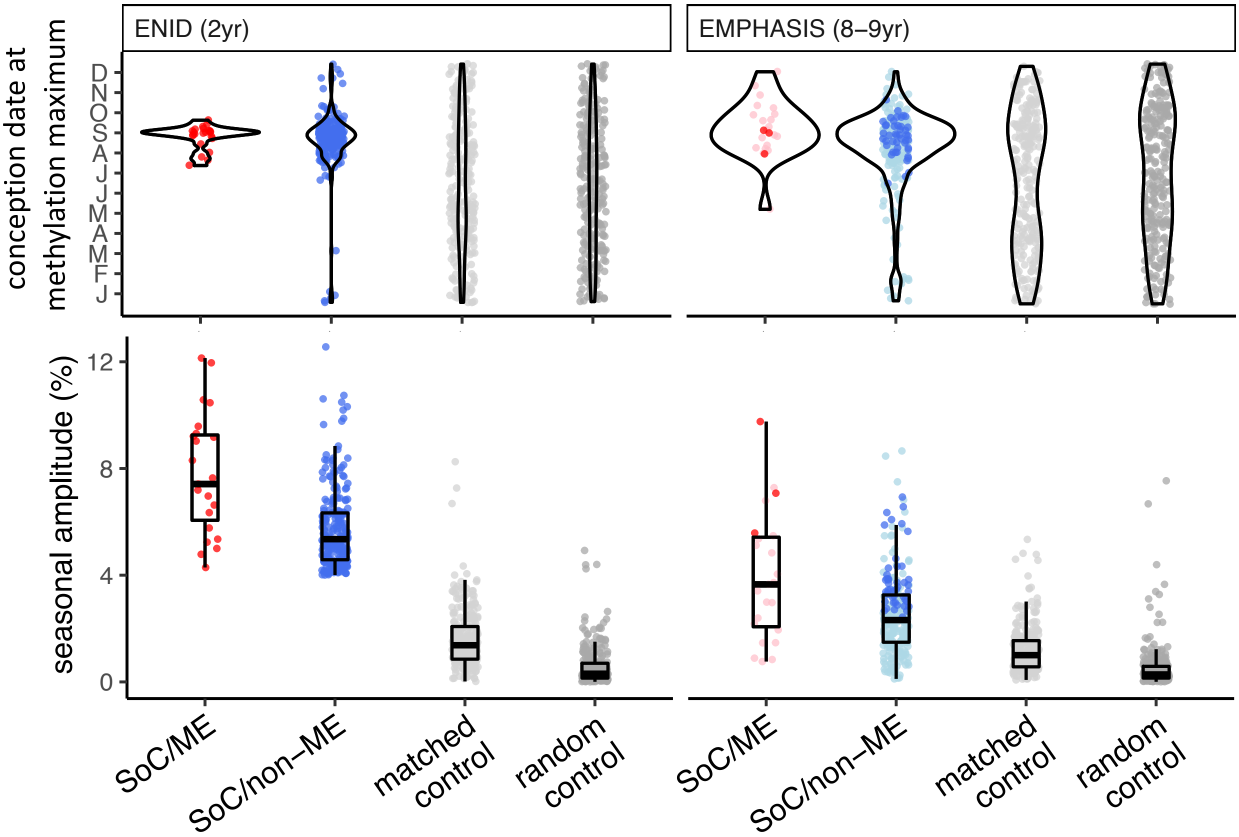


**Supplementary Figure 13. Season of conception analysis adjusted for ethnicity.** This figure replicates Fig. 3C in the main paper, but with additional adjustment for ethnicity. Here we adjust for ethnicity in the EMPHASIS cohort using the first two genetic principle components as covariates in the Fourier regression models (Supplementary Fig. 13A). For the ENID cohort where we do not have genetic data, we adjust for ethnicity using an additional covariate dichotomised according to whether an individual was one of the 9 who came from the predominantly Fula 'outlier village' identified from the PCA using genetic data in the EMPHASIS cohort (Supplementary Fig. 13B).

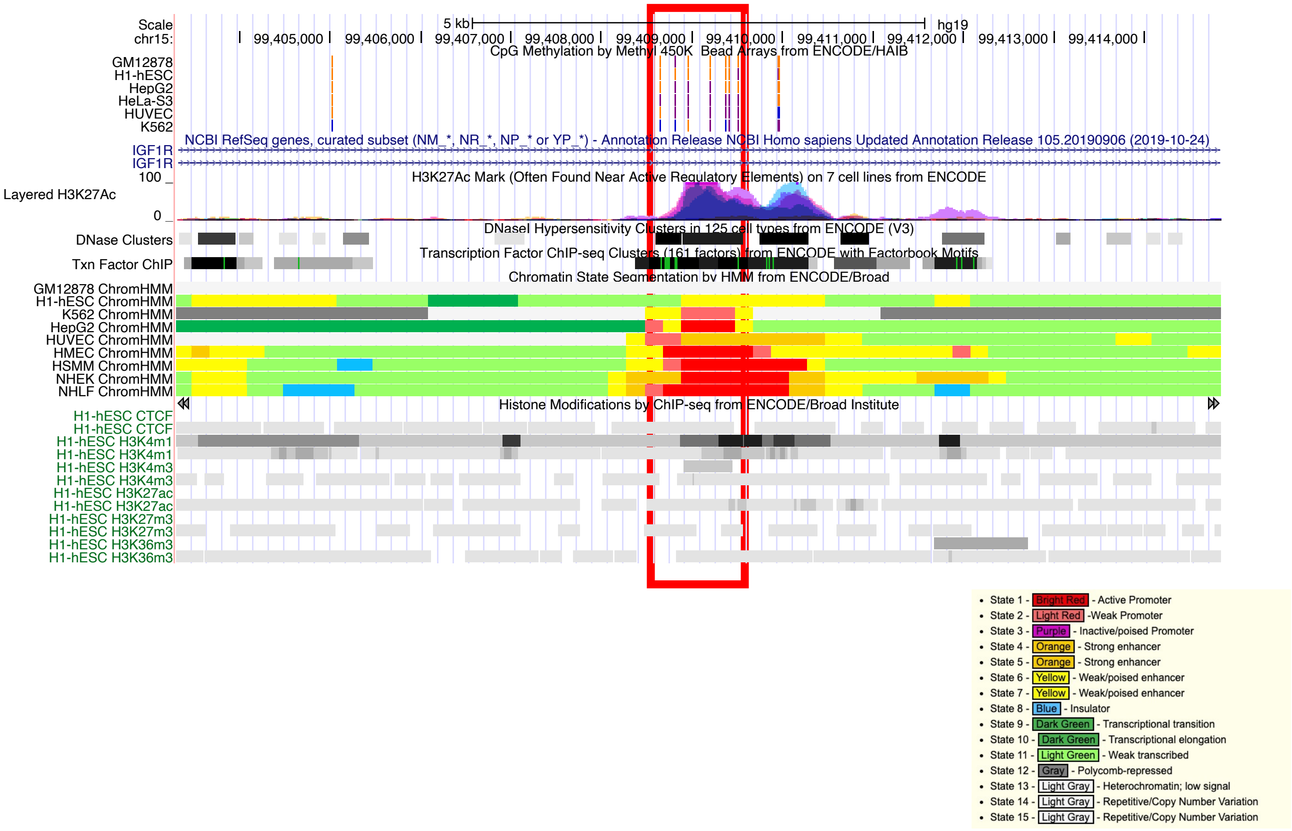


**Supplementary Figure 14. UCSC genome browser plot of the *IGF1R* SoC-CpG cluster.** 7 SoC-CpGs on the Illumina 450k array are highlighted. This region falls within intron 2 and bears the hallmarks of being a promoter and/or active or poised enhancer in multiple cell lines. The key lists colour coding for predicted regulatory regions using ChromHMM.

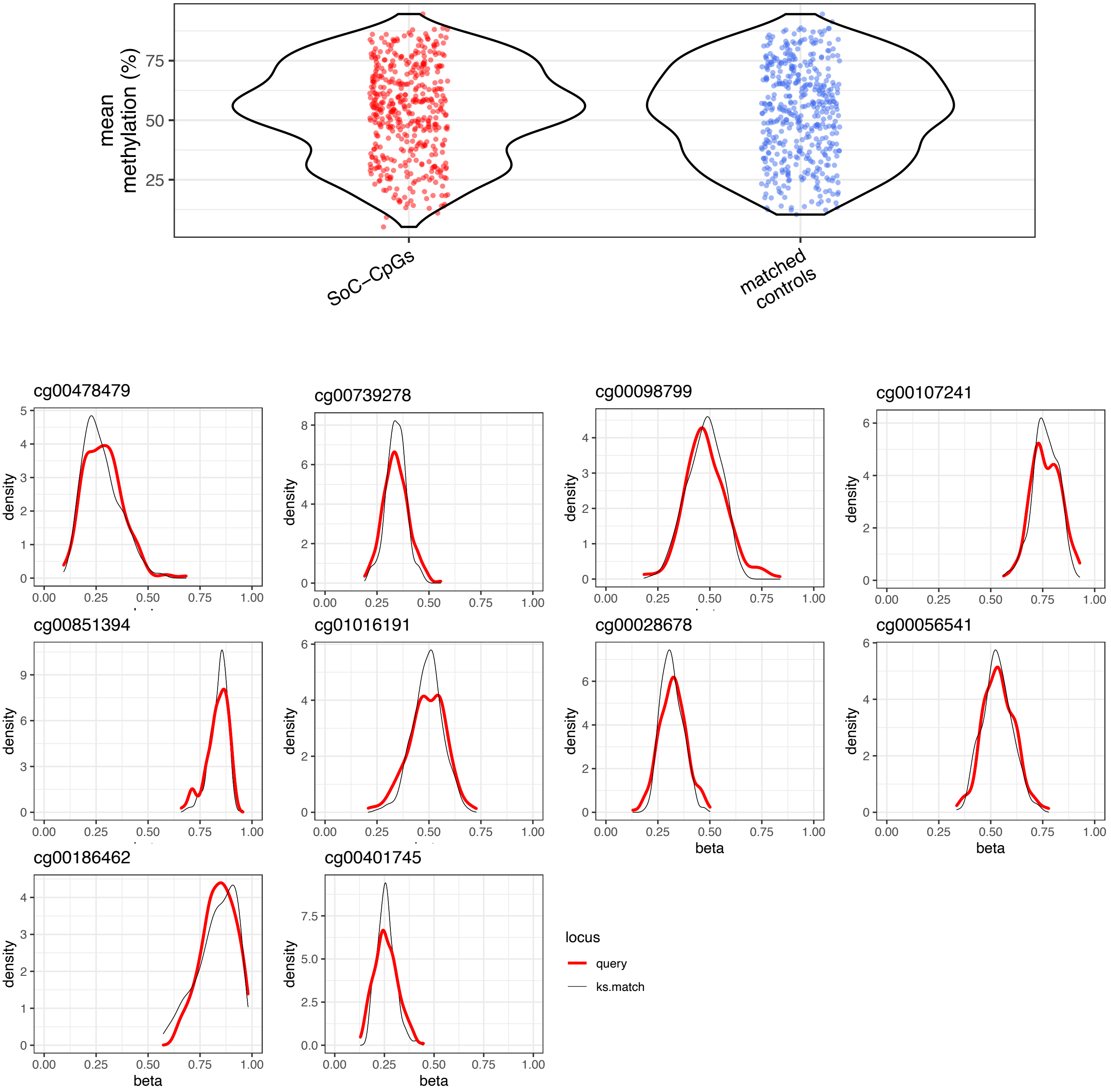


**Supplementary Figure 15. Selection of matched controls.** Matched controls are selected from array background using Kolmogorov-Smirnov (KS) tests to identify CpGs with similar methylation distributions to SoC-CpGs (see Methods). **Top:** Methylation mean distribution at 259 SoC-CpGs (left) vs 259 matched controls (right). **Bottom:** Methylation distribution of 10 SoC-CpGs selected at random (thick red lines) and their corresponding KS-matched controls (thin black lines).

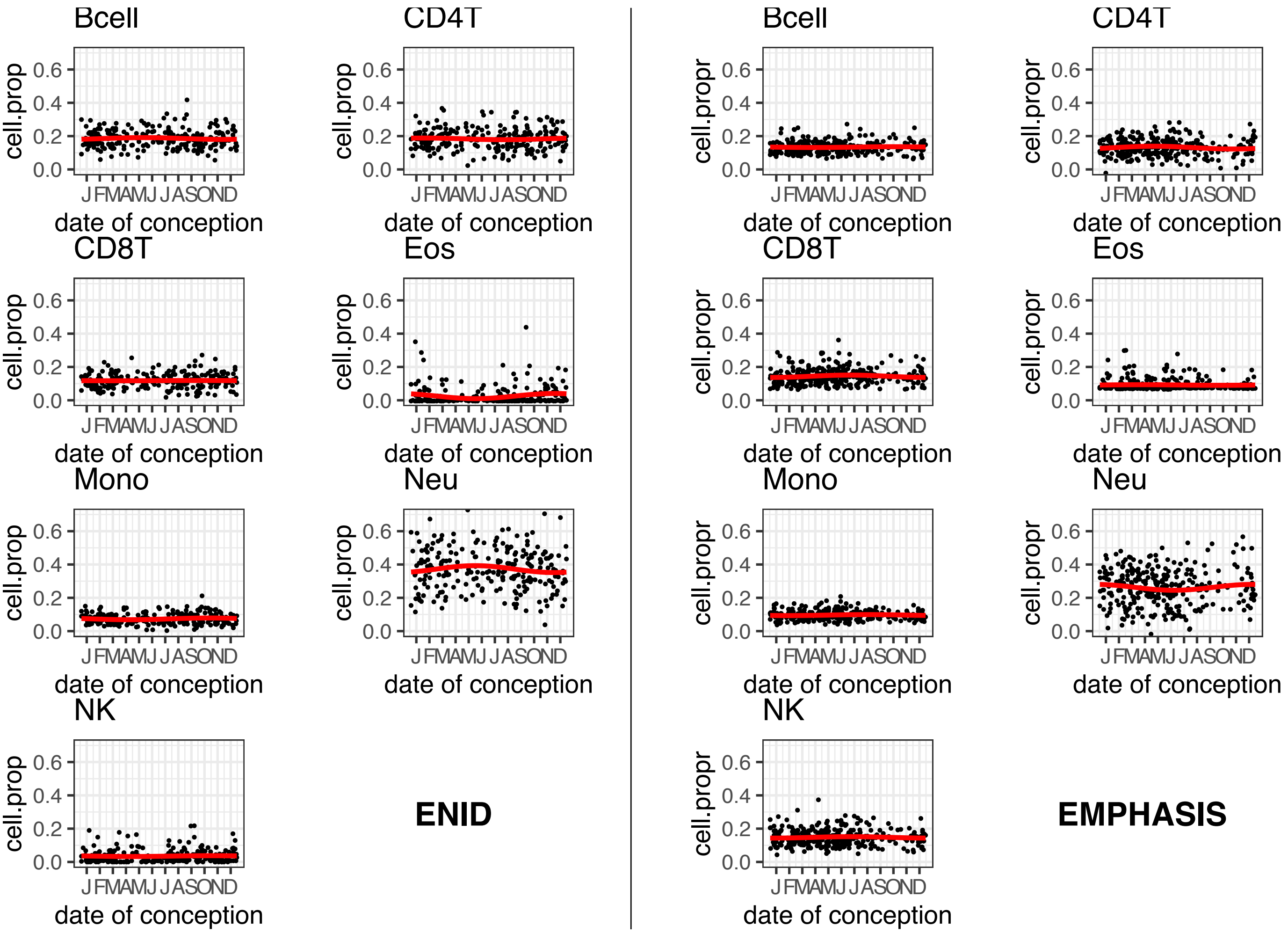


**Supplementary Figure 16. Seasonal variation in estimated cell composition.** Cell proportions for all samples from each cohort are estimated using the estimateCellCounts function from the *minfi* package in R. Seasonal variation (red curves) for each cell type is determined from Fourier regression models adjusted for sex (ENID) or age and sex (EMPHASIS). See Methods for further details.
